# *Pseudomonas aeruginosa* metabolite 3-oxo-C12HSL induces apoptosis through T2R14 and the mitochondrial calcium uniporter

**DOI:** 10.1101/2024.10.24.620094

**Authors:** Zoey A. Miller, Arielle Mueller, Joel C. Thompson, Sarah M. Sywanycz, Brianna L. Hill, Ryan M. Carey, Robert J. Lee

**Affiliations:** Department of Otorhinolaryngology, University of Pennsylvania Perelman School of Medicine, Philadelphia, PA, 19104, USA; Pharmacology Graduate Group, University of Pennsylvania Perelman School of Medicine, Philadelphia, PA, 19104, USA; Department of Physiology, University of Pennsylvania Perelman School of Medicine, Philadelphia, PA, 19104, USA

## Abstract

Head and neck squamous cell carcinomas (HNSCCs) arise in the mucosal lining of the upper aerodigestive tract. HNSCCs have high mortality rates and current treatments can be associated with severe morbidities. It is vital to discover effective, minimally invasive therapies that improve survival and quality of life. We previously discovered that bitter taste receptor 14 (T2R14), a GPCR, kills HNSCC cells when activated by bitter agonists. We are now investigating endogenous bitter ligands that exist in HNSCC tumor microenvironment (TME). The TME includes cells, signaling molecules, and microbes that can greatly influence treatment responses and overall prognosis in HNSCC. *Pseudomonas aeruginosa* is a gram-negative bacterium that colonizes/infects HNSCC patients. 3-oxo-C12SHL is a quorum-sensing N-acyl homoserine lactone (AHL) secreted by *P. aeruginosa* which is also a bitter compound. 3-oxo-C12HSL induces apoptosis but this has never been linked to T2R activation. We hypothesized that 3-oxo-C12HSL induces apoptosis in HNSCC via T2R14. We show that 3-oxo-C12HSL activates intracellular Ca^2+^ responses in HNSCC cells. This is inhibited with T2R14 antagonization. 3-oxo-C12HSL may activate additional Ca^2+^ channels as the Ca^2+^ dynamics are independent from store-operated calcium entry (SOCE). 3-oxo-C12HSL inhibits cell viability, depolarizes mitochondria, and produces ROS. This induces apoptosis in HNSCC cells. In a comparative screen of quorum-sensing AHLs, 3-oxo-C12HSL was the only AHL that elicited both a Ca^2+^ response and reduced cell viability. These results suggest that *P. aeruginosa* may play a significant role in modulating an anti-tumor TME through 3-oxo-C12HSL. Moreover, 3-oxo-C12HSL could be a novel, higher-affinity bitter therapeutic for HNSCC. Further research is warranted to elucidate the mechanisms of other endogenous T2R agonists present in the TME.

## INTRODUCTION

Head and neck squamous cell carcinoma (HNSCC) is the sixth most diagnosed cancer worldwide and the global incidence continues to rise [1, 2]. These cancers arise in the epithelium of the sinonasal tract, oral cavity, larynx, and pharynx [3]. The current five-year survival rate is low at about 50% [1]. These poor outcomes are largely attributed to a lack of preventive screening, late-stage diagnoses, and high rates of metastasis [4]. While surgery, radiation, systemic chemotherapy, and occasionally immunotherapy can be effective, these treatments are often aggressive and invasive [5]. Consequently, patients frequently experience a significant decline in quality of life (QoL) [6]. Side effects of treatment can include loss of verbal communication, physical disfigurement, chronic pain, dependence on feeding tubes and more [7, 8]. Therefore, it is crucial to discover new and effective therapeutics to improve patient QoL during treatment.

We previously uncovered that bitter taste receptors (Taste Family 2 Receptors; T2Rs) could serve as therapeutic targets in HNSCCs [9, 10]. T2Rs are Ca^2+^-activating G protein-coupled receptors (GPCRs) that were first discovered on the tongue but are expressed widely throughout the human body [11]. Our findings showed that HNSCC cells express several T2R isoforms [9, 10]. When T2Rs are activated, HNSCC cells undergo Ca^2+^-dependent apoptosis, driven by mitochondrial Ca^2+^ overload [10]. Agonists for the T2R isoform T2R14 induce robust apoptotic responses [9, 10]. T2R14 is upregulated in some HNSCCs, suggesting that it could be a target for treatment [9].

Our discovery has prompted a new question: do T2Rs have endogenous ligands in the HNSCC tumor microenvironment (TME)? Tumors are surrounded by a network of different cell types, cytokines/chemokines, vasculature, microbes, and metabolites in the TME [12]. These components can facilitate a pro- or anti-tumor environment, thus dictating the prognosis and outcomes of patients [13]. Microorganisms and their metabolites play a significant role in this balance [14]. Interestingly, many gram-negative bacteria, which can colonize the TME, secrete bitter bacterial metabolites, like acyl homoserine lactones (AHLs) and quinolones [15, 16]. T2Rs are known to play diverse roles in innate immunity by sensing these AHLs and quinolones [17–20]. Although there is no evidence to indicate that T2Rs act as immune modulators in HNSCCs, they do serve as pro-apoptotic receptors [9]. We hypothesize that T2Rs could be activated by endogenous bitter bacterial metabolites in the HNSCC TME that may regulate the susceptibility of tumor cells to other death signals.

*Pseudomonas aeruginosa* is a gram-negative bacteria species that can exist in the upper aerodigestive tract as a commensal or opportunistic pathogen [21–23]. *P. aeruginosa* often colonizes or infects HNSCC patients [24]. N-3-oxo-dodecanoyl-L-acylhomoserine lactone (3-oxo-C12HSL) is a quorum-sensing molecule secreted by these bacteria and activates apoptosis in a Ca^2+^-dependent manner [25–28]. *P. aeruginosa* has tumor suppressing properties in several cancers, but the role of 3-oxo-C12HSL in this process has not been explored [29, 30]. The presence of *P. aeruginosa* in tumors is even correlated with positive prognosis [31]

We previously found that 3-oxo-C12HSL is a bitter agonist for T2R38 in the human sinonasal epithelium [32]. In addition, 3-oxo-C12HSL activates heterologously-expressed T2R4, T2R14, and T2R20 [19]. However, the activation of endogenous T2R14 in HNSCCs is unknown. Therefore, we hypothesized that 3-oxo-C12HSL induces apoptosis in HNSCC cells by activating endogenous T2R14.

Here, we show that 3-oxo-C12HSL activates an intracellular Ca^2+^ response through T2R14 in HNSCC cells. 3-oxo-C12HSL decreases cell viability and mitochondrial health, leading to T2R14-mediated apoptosis. This reveals a novel mechanism behind 3-oxo-C12HSL-mediated apoptosis and provides insight on the role that *P. aeruginosa* may play in the HNSCC TME.

## RESULTS

### 3-oxo-C12HSL Activates an Intracellular Ca^2+^ Response in HNSCC Cells

The *TAS2R14* gene encoding T2R14 is the highest expressed *TAS2R* gene across all cancer types in the TCGA [10]. *P. aeruginosa* 3-oxo-C12HSL can activate Ca^2+^ responses [25] and can activate heterologously-expressed T2R14 [19]. We thus hypothesized that 3-oxo-C12HSL mobilizes Ca^2+^ through T2R14 activation in HNSCC cells. To test this, Ca^2+^ responses were recorded in HNSCC cells loaded with Fluo-4 [33]. Significant increases in Ca^2+^ were observed with 100 µM in SCC47 (oral), FaDu (oropharyngeal) and RPMI2650 (nasal) cells (Figure 1A-F; Supplemental Figure 1A). To test if *P. aeruginosa* conditioned media would induce a similar Ca^2+^ response, conditioned LB media from cultured bacteria was collected. Conditioned media from WT strain PAO 1 induced a significant Ca^2+^ response (Figure 1G) while media from strain PAO-JP2 did not induce a Ca^2+^ response (Figure 1G). PAO-JP2 does not produce 3-oxo-C12HSL due to loss of functional *lasI and rhlI* genes [34].

**Figure 1.**
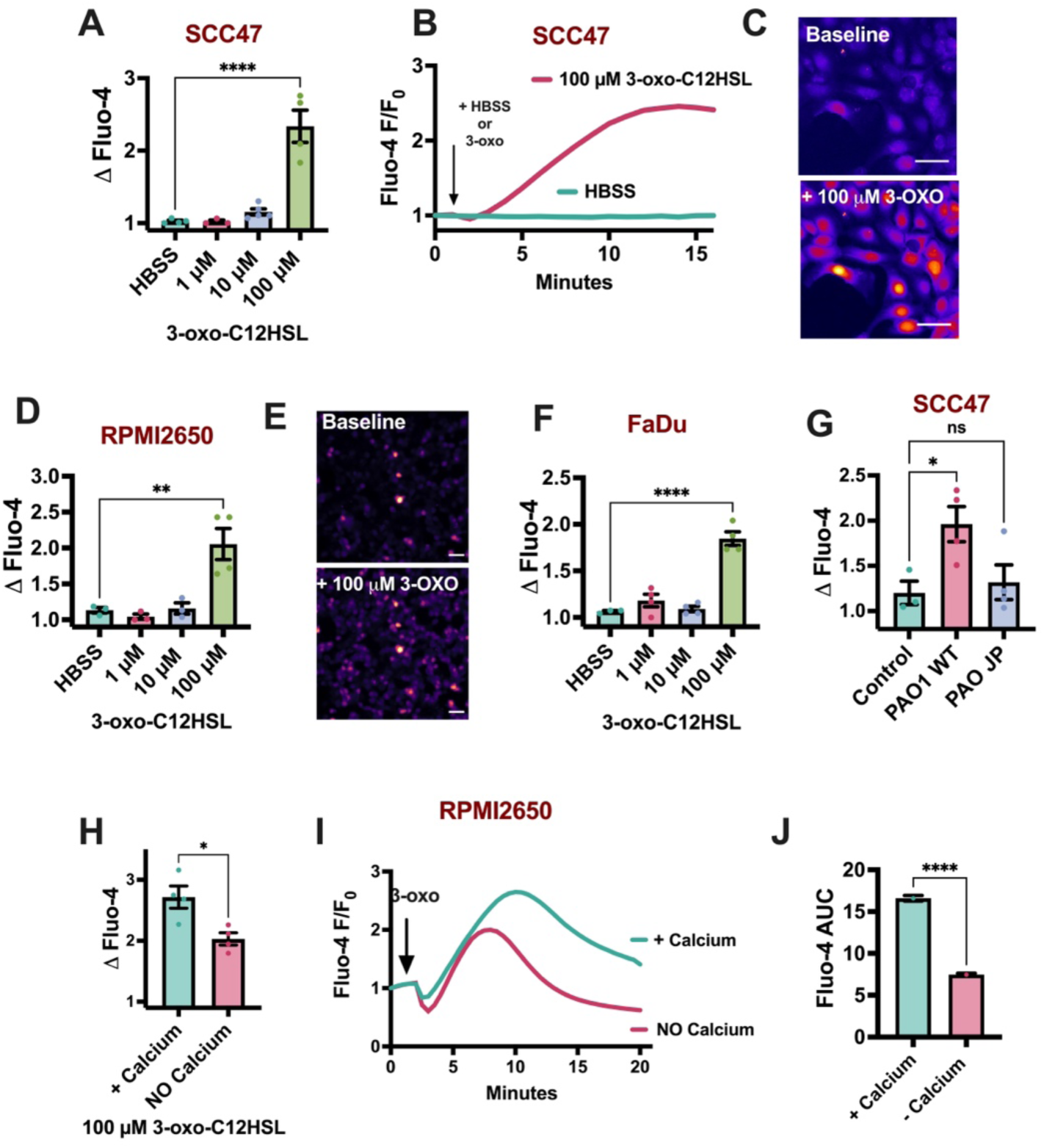
3-oxo-C12HSL mobilizes Ca^2+^ in HNSCC cells. HNSSC cell lines SCC47, FaDu, RPMI2650 were loaded with Fluo-4 and were imaged for Ca^2+^ responses with 3-oxo-C12HSL **A-C)** SCC47 peak Ca^2+^ responses with 0 – 100 µM 3-oxo-C12 (*A*) and Ca^2+^ response over time (*B*) with 100 µM 3-oxo-C12HSL with representative images of baseline and peak Ca^2+^ response (*C*). **D & E)** RPMI2650 peak Ca^2+^ responses with 0 – 100 µM 3-oxo-C12 (*D*) and representative images of baseline and peak Ca^2+^ response (*E*). **F)** FaDu peak Ca^2+^ responses with 0 – 100 µM 3-oxo-C12HSL. **G)** SCC47 peak Ca^2+^ responses with 12.5% conditioned LB media from cultures of PAO1 (WT *P. aeruginosa*) or PAO JP2 (τι*lasI,* τι*rhlI*). Control was HBSS with 12.5% LB, which was highest concentration that did not evoke a significant Ca^2+^ response. See methods for more details. **H-J)** RPMI2650 peak Ca^2+^ responses with 100 µM 3-oxo-C12HSL in HBSS with or without Ca^2+^ (2 mM EGTA) (*H*) and Ca^2+^ response over time (*I*). Area under the curve (AUC) was quantified from response trace (*J*). All traces are representative. All peak Ca^2+^ responses mean ± SEM with >3 experiments using separate cultures. Scale bars = 50 µm. Significance by 1-way ANOVA with Bonferroni’s posttest comparing HBSS/Control to each dose response. Significance by unpaired t-test for +/- Ca^2+^ experiment. P < 0.05 (*), P < 0.01 (**), P < 0.001 (***), and no statistical significance (ns or unmarked).

To further investigate the origin of the Ca^2+^ response, Ca^2+^ imaging was performed with (1.8 mM Ca^2+^) or without (no added Ca^2+^ plus 2 mM EGTA) extracellular Ca^2+^. RPMI2650 cells had a slightly lower but largely intact Ca^2+^ response with 3-oxo-C12HSL in 2 mM EGTA (Figure 1H-J), indicating that a majority of the Ca^2+^ response originates intracellularly [35]. In addition, intracellular Ca^2+^ chelation with BAPTA significantly decreased the Ca^2+^ response with 3-oxo-C12HSL (Supplemental Figure 1B & C). We previously observed larger increases in nuclear Ca^2+^ vs cytoplasmic Ca^2+^ with T2R activation [9, 36]. To measure these specific Ca^2+^ domains, pCMV NES-R-GECO (nuclear export sequence) or pCMV NLS-R-GECO (nuclear localized sequence) were used [37]. 3-oxo-C12HSL increased both cytoplasmic and nuclear Ca^2+^ (Figures 2A-D), with a higher Ca^2+^ response in the nucleus (Figure 2E).

**Figure 2.**
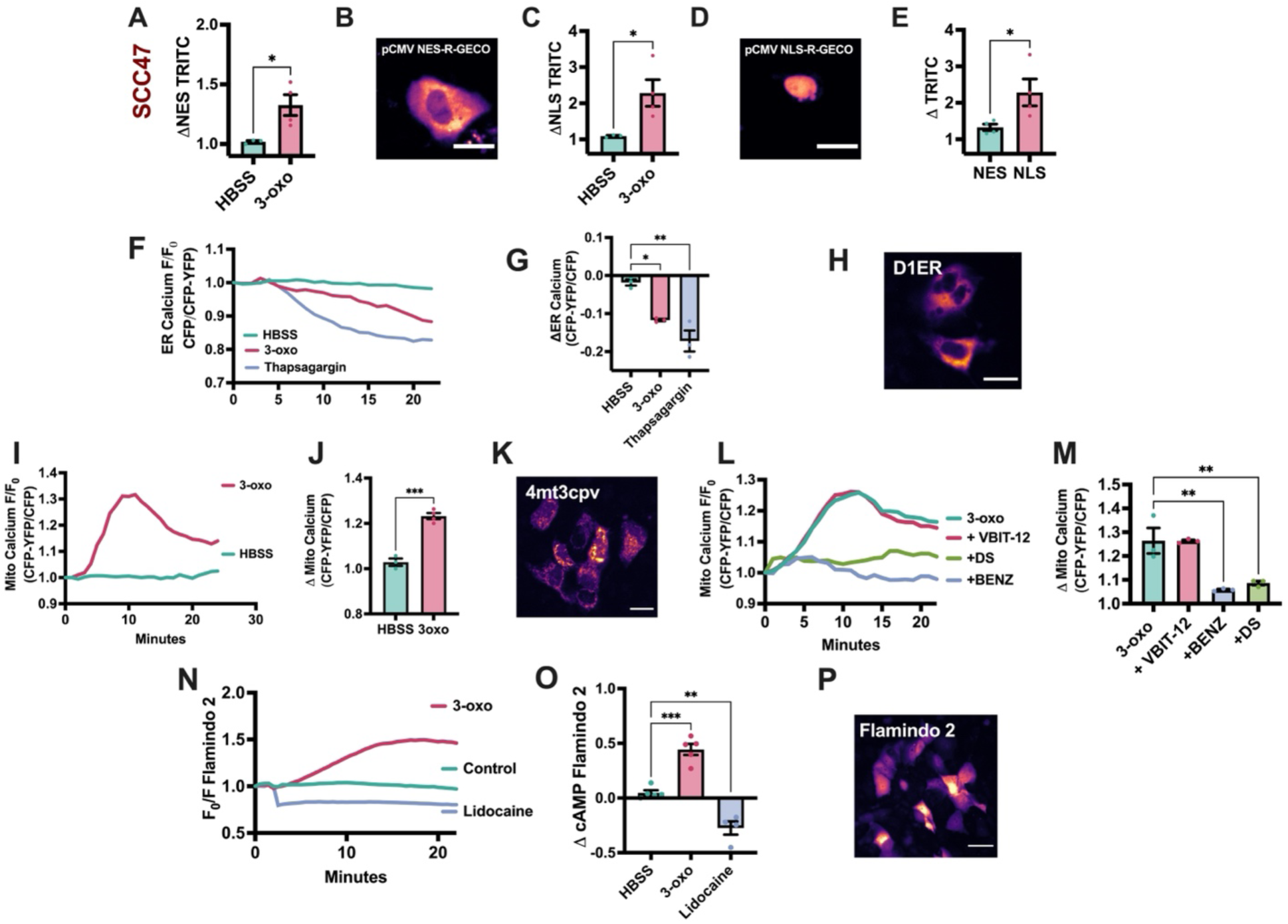
3-oxo-C12HSL activates an intracellular Ca^2+^ response in HNSCC cells. HNSCC cell line SCC47 were loaded with Fluo-4 or transfected with a genetically encoded Ca^2+^ reporter. **A-E)** SCC47 peak cytoplasmic Ca^2+^ response (*A*) reported with pCMV NES-R-GECO (*B*) (scale bar = 20 µm) or peak nuclear Ca^2+^ response (*C*) reported with pCMV NLS-R-GECO (*D*) (scale bar = 20 µm) with 100 µM 3-oxo-C12HSL (3-oxo). Peak cytoplasmic and nuclear Ca^2+^ responses were compared (*E*). **F-H)** SCC47 ER Ca^2+^ efflux over time (*F*) and change (Δ) in CFP-YFP/CFP fluorescence (*G*) with 100 µM 3-oxo-C12HSL (3-oxo) or 10 µg/mL thapsigargin (Thapsigargin) using D1ER Ca^2+^ reporter (*H*) (scale bar = 30 µm). **I-K)** SCC47 mitochondrial Ca^2+^ influx over time (*I*) and change (Δ) in CFP-YFP/CFP fluorescence (J) with 100 µM 3-oxo-C12HSL (3-oxo) using 4mt3Dcpv mitochondrial Ca^2+^ reporter (*K*) (scale bar = 30 µm). **L & M)** SCC47 mitochondrial Ca^2+^ influx over time with (L) and change (Δ) in CFP-YFP/CFP fluorescence (*M*) with 100 µM 3-oxo-C12HSL (3-oxo) +/- 50 µM VBIT-12 (VBIT-12), 50 µM DS16570511 (DS), or 50 µM benzethonium chloride (BENZ). **N-P)** SCC47 cAMP response over time (*N*) and change (Δ) FITC fluorescence O*P*) with 100 µM 3-oxo-C12HSL (3-oxo) or 5 mM lidocaine (Lidocaine) using cAMP reporter Flamindo 2 (*P*) (scale bar = 30 µm). All traces are representative. All bar graphs mean ± SEM with >3 experiments using separate cultures. Significance by 1-way ANOVA with Dunnett posttest comparing HBSS/Control to each dose response when comparing more than two conditions. Significance by unpaired t-test when testing two conditions. P < 0.05 (*), P < 0.01 (**), P < 0.001 (***), and no statistical significance (ns or unmarked).

T2Rs typically stimulate Ca^2+^ release from the ER via the inositol trisphosphate (IP_3_) receptor (IP_3_R), with sustained Ca^2+^ responses requiring store-operated Ca^2+^ entry (SOCE) [9, 10, 36]. An ER-targeted biosensor, D1ER, was used to measure ER Ca^2+^ dynamics [38]. 3-oxo-C12-HSL significantly decreased ER Ca^2+^ over 20 minutes (Figure 2F-H). These results were confirmed using a different ER-localized Ca^2+^ biosensor, ER-LAR GECO (Supplemental Figure 1D-F) [39]. These data support activation of a GPCR pathway, possibly a T2R, by 3-oxo-C12HSL. T2R activation also causes an influx of Ca^2+^ into the mitochondria [40, 41]. Using mitochondrial Ca^2+^ biosensor 4mt3cpv [38, 39], we saw that 3-oxo-C12-HSL increased mitochondrial Ca^2+^ (Figure 2I-K). This was inhibited when the mitochondrial calcium uniporter (MCU), a Ca^2+^ channel on the inner membrane of the mitochondria, was inhibited with benzethonium chloride or DS16570511 (Figure 2L & M) [42–44]. Mitochondrial Ca^2+^ was unchanged in the presence of VBIT-12, a purported inhibitor of voltage-dependent anion channel 1 (VDAC1) (Figure 2K-M) [45]. VDAC channels conduct Ca^2+^ across the outer membrane of the mitochondria, and while VDAC1 is the highest expressed isoform in HNSCC cells (Supplemental Figure 1G) [42], VBIT-12 had no effect on mitochondrial Ca^2+^ increases either with 3-oxo-C12HSL or ATP, which activates Ca^2+^ via purinergic receptors (Supplemental Figure 1H-K) [46].

To further investigate T2R-specific downstream signaling, we used EPAC-based cAMP biosensor Flamindo2 to measure intracellular cAMP dynamics [47]. T2R activation of Gαi inhibits adenylyl cyclase, decreasing cAMP [48]. Surprisingly, 3-oxo-C12HSL increased cAMP (Figure 2N-P). Lidocaine (T2R14 agonist) and β-adrenergic agonist isoproterenol were used as positive controls (Figure 2N & O; Supplemental Figure 2A & B). To test if the increase in cAMP with 3-oxo-C12HSL was Ca^2+^-dependent, intracellular Ca^2+^ was chelated with BAPTA as done for Ca^2+^ experiments above. The 3-oxo-C12HSL response was unchanged (Supplemental Figure 2C). 3-oxo-C12HSL also activated cAMP-dependent protein kinase A, observed with Sapphire-AKAR biosensor (Supplemental Figure 2D-F) [49, 50]. Despite links between Ca^2+^ and activation of AMP-dependent protein kinase (AMPK), we also observed that 3-oxo-C12HSL did not activate AMPK activity using the AMPKAR biosensor, while we did see significant increase in AMPK activity with ionomycin (Supplemental Figure 2G & H) [51].

### Sustained 3-oxo-C12HSL Ca^2+^ Mobilization Is Independent from Store-Operated Calcium Entry

T2R activation typically causes an efflux of Ca^2+^ from the ER [10], supported by our data above. Sustained Ca^2+^ signaling requires SOCE [52]. When ER Ca^2+^ stores are depleted, ER-resident STIM proteins sense the loss of ER Ca^2+^ content and activate plasma membrane Orai Ca^2+^ channels to allow Ca^2+^ influx [53]. To test if the Ca^2+^ response from 3-oxo-C12HSL utilizes SOCE, SCC47 and RPMI2650 cells were pre-treated with BPT2, an inhibitor of ORAI isoform 1 (ORAI1) [54]. Cells were first stimulated with 3-oxo-C12HSL in 0 Ca^2+^ HBSS (2 mM EGTA) +/- BTP2. After the initial response, 2 mM Ca^2+^ was added-back to stimulate Ca^2+^ influx via SOCE in the continued presence or absence of BTP2. Surprisingly, cells with and without BTP2 both exhibited Ca^2+^ influx with Ca^2+^ addback (Figure 3A-D; Supplemental Figure 3A). When this experiment was repeated using thapsagargin, a sarco/endoplasmic reticulum Ca^2+^ ATPase (SERCA) pump inhibitor [55], BPT2 blocked Ca^2+^ influx as expected (Figure 3E-H).

**Figure 3.**
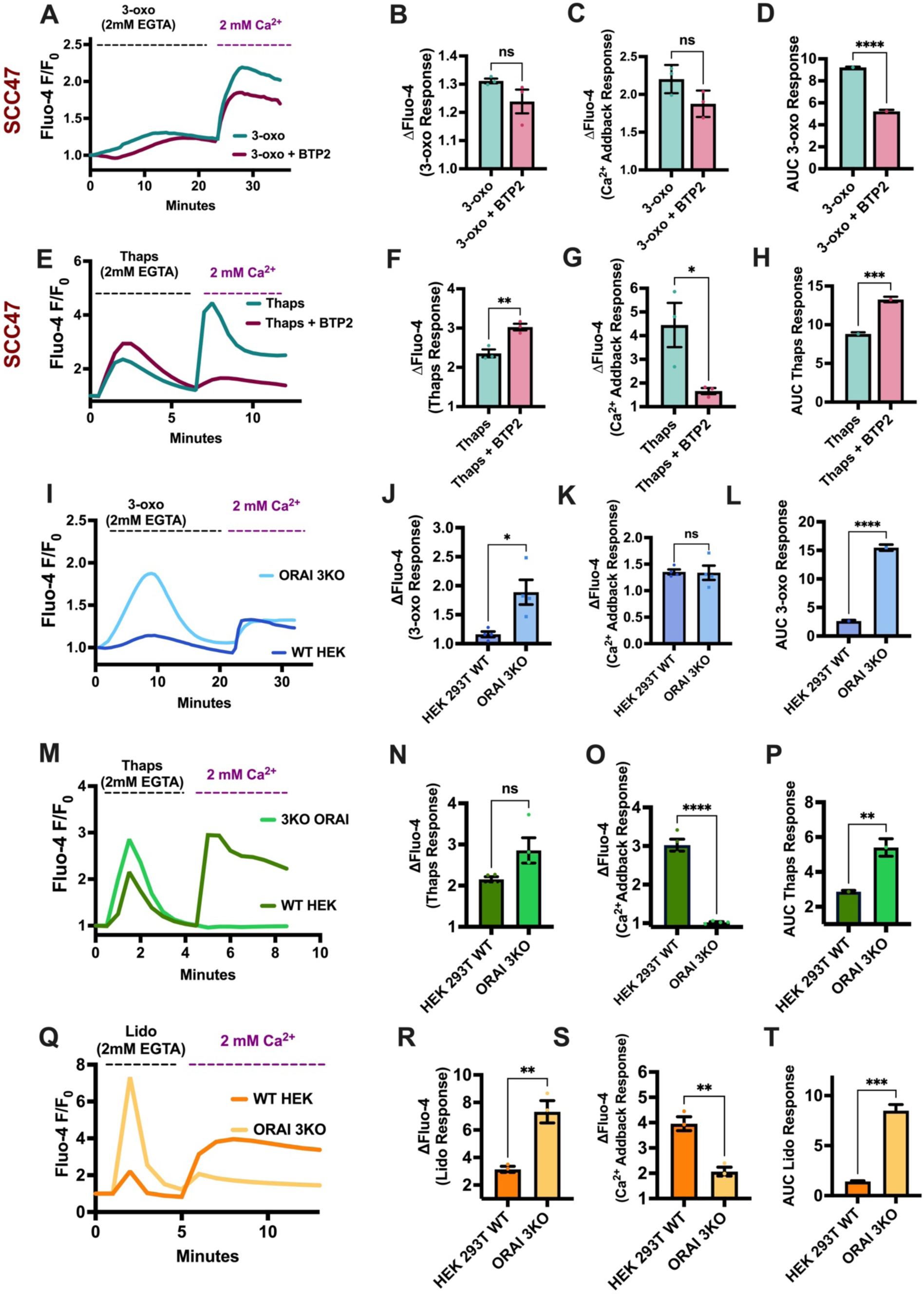
Sustained 3-oxo-C12HSL Ca^2+^ mobilization is independent from Orai-Stim store-operated calcium entry. **A-D)** SCC47 Ca^2+^ trace (*A*) and peak responses to first stimulation with 100 µM 3-oxo-C12SHL (3-oxo) in 2 mM EGTA (*B*) and subsequent stimulation with 2 mM Ca^2+^ addback (*C) +/-* 10 µM BTP2. Area under the curve (AUC) was quantified from primary Ca^2+^ trace with 3-oxo *(D*). **E-H)** SCC47 Ca^2+^ trace (E) and peak responses to first stimulation with 100 µM 3-oxo-C12SHL (3-oxo) in 2 mM EGTA (*F*) and subsequent stimulation with 2 mM Ca^2+^ addback (*G)* 10 µM BTP2. Area under the curve (AUC) was quantified from primary Ca^2+^ trace with 3-oxo *(H*). **I-L)** WT and ORAI 3KO HEK Ca^2+^ trace (*I*) and peak responses to first stimulation with 100 µM 3-oxo-C12SHL (3-oxo) in 2 mM EGTA (*J*) and subsequent stimulation with 2 mM Ca^2+^ addback (*K*). Area under the curve (AUC) was quantified from primary Ca^2+^ trace with 3-oxo (*L*). **M-P**) WT HEK 293T and ORAI 3KO HEK cells Ca^2+^ response over time (*M*) and peak responses to primary stimulation with 10 µM thapsigargin (Thaps) in 2 mM EGTA (*N*) and secondary stimulation with 2 mM Ca^2+^ addback (*O*). Area under the curve (AUC) was quantified from primary Ca^2+^ trace with Thaps (*P*). **Q-T)** WT HEK 293T and ORAI 3KO HEK cells Ca^2+^ response over time (*Q*) and peak responses to primary stimulation with 10 mM lidocaine (Lido) in 2 mM EGTA (*R*) and secondary stimulation with 2 mM Ca^2+^ addback (*S*). Area under the curve (AUC) was quantified from primary Ca^2+^ trace with Lido (*T*). All traces are representative. All bar graphs mean ± SEM with >3 experiments using separate cultures. Significance by unpaired t-test +/- Ca^2+^ experiment when testing two conditions. P < 0.05 (*), P < 0.01 (**), P < 0.001 (***), and no statistical significance (ns or unmarked).

To further test for SOCE in the 3-oxo-C12HSL Ca^2+^ response, Ca^2+^ addback responses post-agonist stimulation were measured in ORAI Stable Triple Knockout HEK293Ts (ORAI 3KO). As with BTP2, 3-oxo-C12HSL-stimulated ORAI 3KO cells still exhibited Ca^2+^ add-back responses (Figure 3I-L). Interestingly, 3-oxo-C12HSL activated larger Ca^2+^ response in ORAI 3KO than WT HEK293Ts (Figure 3J & L). Unlike 3-oxo-C12HSL, Ca^2+^ add-back responses after thapsagargin or lidocaine stimulation was blocked in ORAI 3KO, as expected (Figure 3M-T). Other similar Ca^2+^ add-back protocols supported STIM-Orai SOCE by T2R14 agonist lidocaine but not 3-oxo-C12HSL (Supplemental Figure 3B-C). Interestingly, as with 3-oxo-C12HSL, lidocaine modulated a higher Ca^2+^ response in ORAI 3KO cells than in HEK293T WT cells (Figure 3R & T). Heterologous expression of ORAI1 in ORAI 3KO cells slightly decreased the Ca^2+^ response, showing that the presence of ORA1 may affect T2R14-induced Ca^2+^ responses. (Supplemental Figure 3C & D). *TAS2R14* mRNA expression was highest in ORAI 3KO cells vs FaDu, RPMI2650, or WT HEK293Ts, which may also contribute to the increased 3-oxo-C12HSL and lidocaine Ca^2+^ responses (Supplemental Figure 3E).

Other Ca^2+^ channels can mediate Ca^2+^ influx into non-excitable cells [56], and some can be activated following GPCR stimulation. Others have hypothesized that 3-oxo-C12HSL activates Ca^2+^-permeable transient receptor potential cation channel subfamily V member 1 (TRPV1) [57]. We observed TRPV1 expression in all cell types utilized in this study (Supplemental Figure 3F) [58]. Interestingly, while SCC47 and ORAI 3KO cells exhibited Ca^2+^ responses to TRPV1 agonist capsaicin [59], HEK293T WT cells did not (Supplemental Figure 3G - J). Further work is needed to determine relationship between Orai and *TAS2R14* expression and TRPV1 function, but our data nonetheless support a lack of activation of Orai-Stim during 3-oxo-C12HSL stimulation, despite activation of ER Ca^2+^ release. This differs from other T2R agonist we have examined.

### 3-oxo-C12HSL-evoked Ca^2+^ responses involve T2R14

To test if 3-oxo-C12HSL activated a ER Ca^2+^ release via phospholipase C (PLC) generation of IP_3_, we used PLC inhibitor U-73122 or inactive analogue U-73343 [60]. Inhibition of PLC significantly decreased the 3-oxo Ca^2+^ response (Figure 4A & B), supporting GPCR involvement. T2R14 antagonists LF1, LF22, or 6-methoxyflavone [10, 61] all significantly dampened the 3-oxo-C12HSL Ca^2+^ response in SCC47, FaDu, and RPMI2650 cells (Figure 4C-K). As described above, culture supernatant from WT PAO1 induced a Ca^2+^ response in SCC47 cells (Figure 1G). Antagonizing T2R14 also significantly decreased the Ca^2+^ response with *P. aeruginosa* PAO1 conditioned media (Figure 4L). The JP2 strain was used as an additional control (Figure 4L). While this supports a pivotal role for T2R14 in the response to *P. aeruginosa* secreted metabolites, it does not specifically implicate 3-oxo-C12HSL, as *P. aeruginosa* also produce N-butyryl-L-homoserine lactone (C4HSL) [62]. In addition, the culture media could contain other unknown Ca^2+^ activating agonists. We previously showed that these antagonists do not modify Ca^2+^ response to many other T2R or purinergic receptor stimuli, so our data nonetheless suggest an important role for T2R14 in the acute Ca^2+^ response to *P. aeruginosa*.

**Figure 4.**
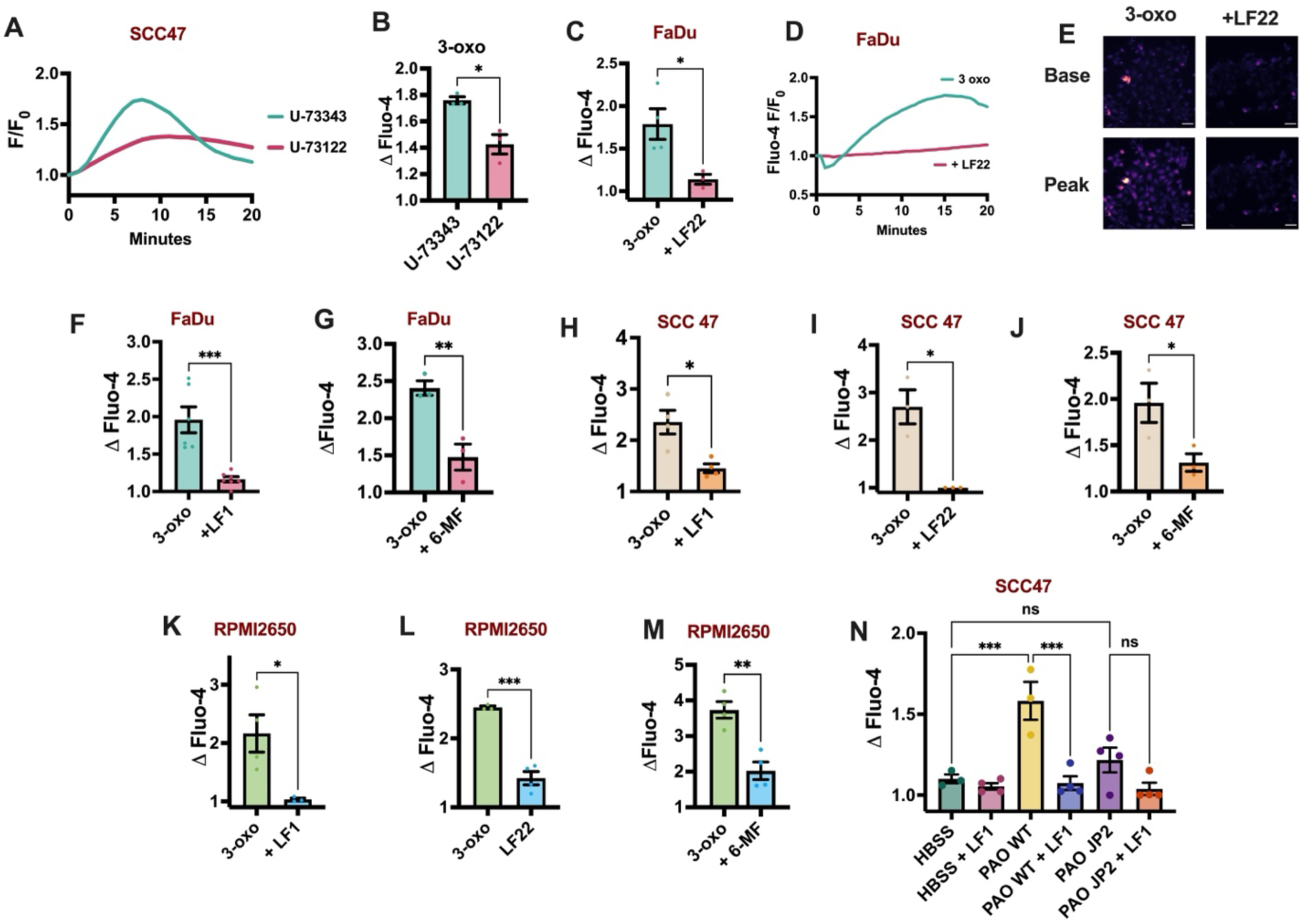
3-oxo-C12HSL activates T2R14 in HNSCC cells. SCC47, FaDu, and RPMI2650 cells were loaded with Fluo-4 and imaged for Ca^2+^ responses. **A & B)** SCC47 Ca^2+^ response over time (*A*) and peak Ca^2+^ response (*B*) with 100 µM 3-oxo-C12SHL (3-oxo) + 1 µM U73343 (negative control analogue) or 1 µM U73122 (PLC inhibitor). **C-E)** FaDu peak Ca^2+^ response (*C*) and Ca^2+^ response over time (*D*) with 100 µM 3-oxo-C12SHL (3-oxo) +/- 500 µM LF22. Representative images shown on the right (*E*). **F & G)** FaDu peak Ca^2+^ response with 100 µM 3-oxo-C12SHL (3-oxo) +/- 500 µM LF21 (*F*) or +/- 100 µM 6-methoxyflavonr (6-MF) (*G*). **H-J)** SCC47 peak Ca^2+^ response with 100 µM 3-oxo-C12SHL (3-oxo) +/- 500 µM LF1 (*H*), 500 µM LF22 (*I*), or 100 µM 6-methoxyflavone (6-MF) (*J*). **K-M)** RPMI2650 peak Ca^2+^ response with 100 µM 3-oxo-C12SHL (3-oxo) +/- 500 µM LF1 (*K*), 500 µM LF22 (*L*), or 100 µM 6-methoxyflavone (6-MF) (*M*) **L)** SCC47 peak Ca^2+^ responses with LB cultured supernatant from PAO1 (WT *P. aeruginosa*) or PAO JP2 (mutated *lasI and rhlI* genes) +/- 500 µM LF1. Control was HBSS with 12.5% LB, which was highest concentration that did not evoke a significant Ca^2+^ response. See methods for more details. All traces are representative. All bar graphs mean ± SEM with >3 experiments using separate cultures. Scale bars = 50 µm. Significance by unpaired t-test. 1-way ANOVA was performed on (*L*) with multiple comparisons (HBSS vs PAO1 WT; HBSS vs PAO JP2; PAO1 WT vs PAO1 WT + LF1; PAO JP2 vs PAO JP2 + LF1) with Šidák correction. P < 0.05 (*), P < 0.01 (**), P < 0.001 (***), and no statistical significance (ns or unmarked).

### 3-oxo-C12HSL Decreases Cell Viability

3-oxo-C12HSL can decrease cell viability and activates apoptosis in a variety of cell types, including some cancer cells [27, 28, 63–66]. We hypothesized that the quorum-sensing molecule might affect cell viability and metabolism through T2R14, which we previously showed activates apoptotic responses in HNSCC cells. Using crystal violet staining, we found that 100 µM 3-oxo-C12HSL decreased cell viability over 24-48 hours in SCC47, FaDu, and RPMI2650 cells (Figure 5A-C; Supplemental Figure 4A-C). Orai inhibitor BPT2 did not enhance or rescue viability effects of 3-oxo-C12HSL (Supplemental Figure 4D), as would be expected due to our data suggesting lack of involvement of Orai. 3-oxo-C12HSL did not affect cell viability in HEK293T WT or ORAI 3KO HEK293 cells (Supplemental Figure 4E). Capsaicin had only a small effect (∼5%) on cell viability (Supplemental Figure 4F), suggesting that activation of TRPV1 is not sufficient to mimic the effects of 3-oxo-C12HSL. To further measure cell viability, we used an XTT assay to indirectly measure NADH production [67]. As low as 10 µM 3-oxo-C12HSL decreased NADH production over 6 hours in SCC47, FaDu, and RPMI2650 cells (Figure 5D-I).

**Figure 5.**
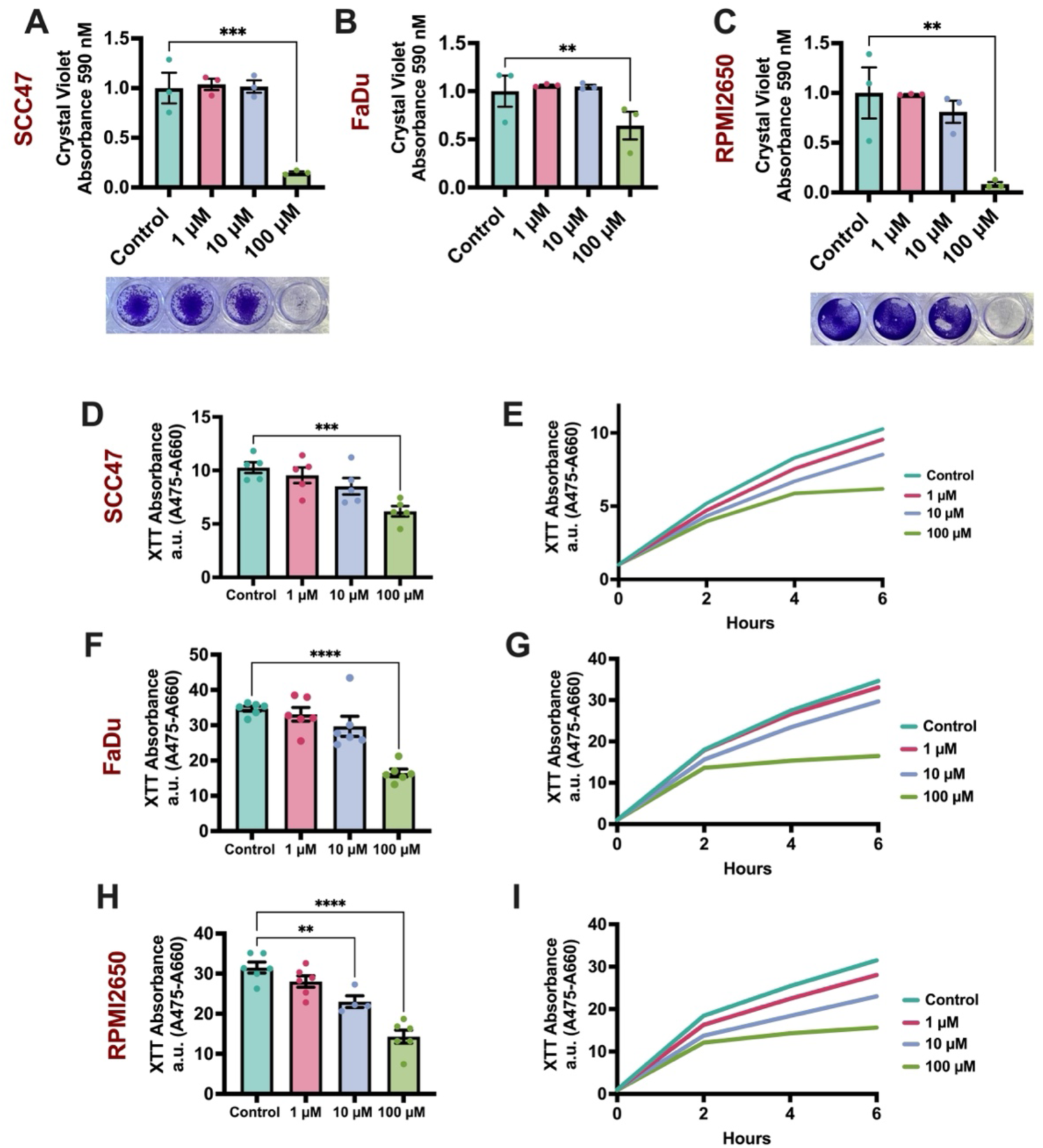
3-oxo-C12HSL decreases cell viability. HNSCC cell viability was measured with crystal violet assay, **A-C)** SCC47 (*A*), FaDu (*B*), and RPMI2650 (*C*) cells were treated with 0 – 100 µM 3-oxo-C12HSL for 24 hours in normal media conditions and stained with crystal violet, indicating alive/adherent cells post-treatment. Solubilized crystal violet absorbance was measured at 590 nM and quantified. **D & E)** SCC47 XTT absorbance (475 nM – 660 nM) value at (*D*) and over (*E*) 6 hours with 0 – 100 µM 3-oxo-C12HSL. **F & G)** FaDu XTT absorbance (475 nM – 660 nM) value at (*F*) and over (*G*) 6 hours with 0 – 100 µM 3-oxo-C12HSL. **H & I)** RPMI2650 absorbance (475 nM – 660 nM) value at (*H*) and over (*I*) 6 hours with 0 – 100 µM 3-oxo-C12HSL. All traces are representative. All bar graphs mean ± SEM with >3 experiments using separate cultures. Significance by 1-way ANOVA with Dunnett posttest comparing 3-oxo-C12HSL treatment to control (media only). P < 0.05 (*), P < 0.01 (**), P < 0.001 (***), and no statistical significance (ns or unmarked).

### Mitochondrial Depolarization with 3-oxo-C12HSL Is Mediated by T2R14 and MCU

We previously found that T2R activation causes mitochondrial Ca^2+^ overload in HNSCC cells [9, 10]. The Ca^2+^ overload causes excessive ROS production and drives mitochondrial-dependent apoptosis [10]. Using TMRE dye, which measures mitochondrial membrane potential, we found that 3-oxo-C12HSL significantly depolarized mitochondrial potential over 15 hours in SCC47 cells (Figure 6A-C) [68]. Additionally, MitoSox, a fluorescent superoxide indicator, showed that 3-oxo-C12HSL also stimulated mitochondrial ROS over 15 hours in SCC47 cells (Figure 6D-F) [10]. 3-oxo-C12HSL had less pronounced effects on mitochondrial membrane potential and ROS in FaDu cells (Supplemental Figure 5A-D) but more strongly affected these parameters in RPMI2650 cells (Supplemental Figure 5E-H). When T2R14 was antagonized with 6-MF, mitochondrial depolarization and ROS production with 3-oxo-C12HSL were rescued in SCC47 (Figure 6G-J) and RPMI2650 (Supplemental Figure 5E-F) cells. Inhibiting MCU inhibited mitochondrial Ca^2+^ uptake (Figure 2K-M), and when MCU was inhibited with benzethonium chloride, the reduction of MMP and ROS with 3-oxo-C12HSL was blocked in SCC47 (Figure 6K-P) and RPMI2650 cells (Supplemental Figure 5G-H). VDAC1 inhibitor VBIT-12 did not alter MMP depolarization with 3-oxo-C12HSL (Supplemental Figure 5I & J).

**Figure 6.**
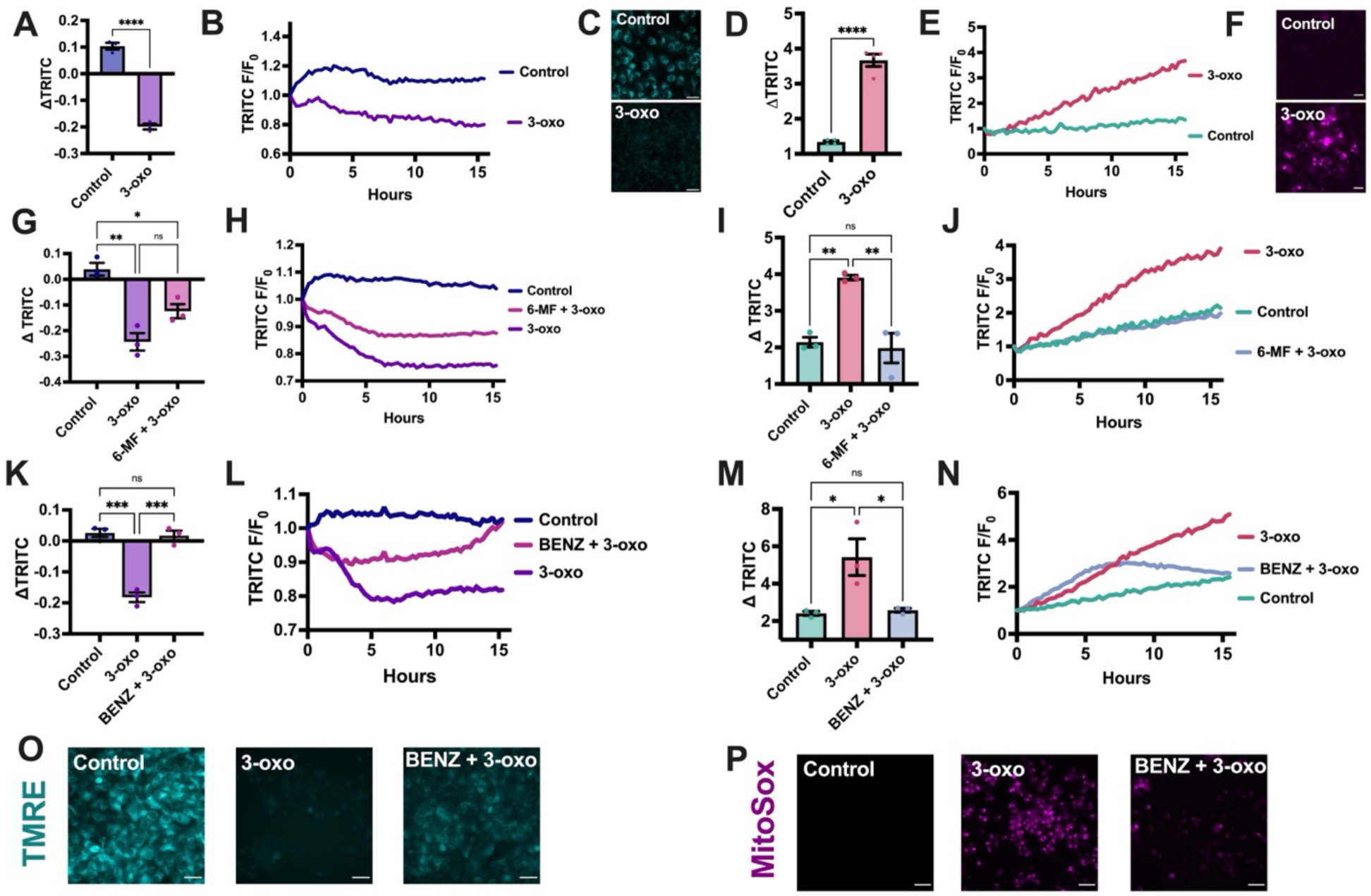
Mitochondrial depolarization with 3-oxo-C12HSL is mediated by T2R14 and MCU. Mitochondrial health and ROS production was measured SCC47 cells. **A-C)** ΔTMRE fluorescence after 15 hours (*A*) and over time (*B*) with 100 µM 3-oxo-C12HSL (3-oxo). Representative images of TMRE fluorescence at 15 hours +/- 3-oxo (*C*). **D-F)** ΔMitoSox fluorescence after 15 hours (*D*) and over time (*E*) with 100 µM 3-oxo-C12HSL (3-oxo). Representative images of MitoSox fluorescence at 15 hours +/- 3-oxo (*F*). **G & H)** ΔTMRE fluorescence after 15 hours (*G*) and over time (*H*) with 100 µM 3-oxo-C12HSL (3-oxo) +/- 100 µM 6-MF. **I & J)** ΔMitoSox fluorescence after 15 hours (*I*) and over time (*J*) with 100 µM 3-oxo-C12HSL (3-oxo) +/- 6-MF. **K & L)** ΔTMRE fluorescence after 15 hours (*K*) and over time (*L*) with 100 µM 3-oxo-C12HSL (3-oxo) +/- 100 µM 6-MF. **M & N)** ΔMitoSox fluorescence after 15 hours (*M*) and over time (*N*) with 100 µM 3-oxo-C12HSL (3-oxo) +/- 50 µM benzethonium chloride (BENZ). **O & P)** Representative images of TMRE (*O*) and MitoSox (*P*) fluorescence after 15 hours with 100 µM 3-oxo-C12HSL (3-oxo) +/- 50 µM benzethonium chloride (BENZ). All traces are representative. All bar graphs mean ± SEM with >3 experiments using separate cultures. Significance for two conditions determined with unpaired t-test. Scale bars = 50 µm. Significance for more than two conditions was determined with 1-way ANOVA with multiple comparisons (Control vs 3-oxo; Control vs 3-oxo + 6-MF or BENZ; 3-oxo vs 3-oxo + 6-MF or BENZ) with Šidák correction. (P < 0.05 (*), P < 0.01 (**), P < 0.001 (***), and no statistical significance (ns or unmarked).

### 3-oxo-C12HSL Induces Apoptosis via T2R14

To test if 3-oxo-C12HSL induced apoptosis in HNSCC cells via T2R14, a fluorescent DEVD peptide assays (CellEvent) was used to measure caspase-3 and −7 cleavage. 3-oxo-C12HSL induced apoptosis over 15 hours in SCC47 and RPMI2650 cells (Figure 7A-E). 3-oxo-C12HSL also inhibited FaDu cell anchorage-independent spheroid formation over 24 hours (Figure 7F). In parallel to our observations with mitochondrial health, apoptosis was blocked when either 6-MF or MCU were inhibited with 6-MF or benzethonium chloride, respectively, in the presence of 3-oxo-C12HSL in both SCC47 and RPMI2650 cells (Figure 7G-L). Capsaicin had no effect on apoptosis in SCC47 cells (Supplemental Figure 6A & B), again supporting that TRPV1 activation is not sufficient for mimicking the effects of 3-oxo-C12HSL. VDAC1 inhibitor VBIT-12 had no effect on apoptosis with 3-oxo-C12HSL (Supplemental Figure 6C & D).

**Figure 7.**
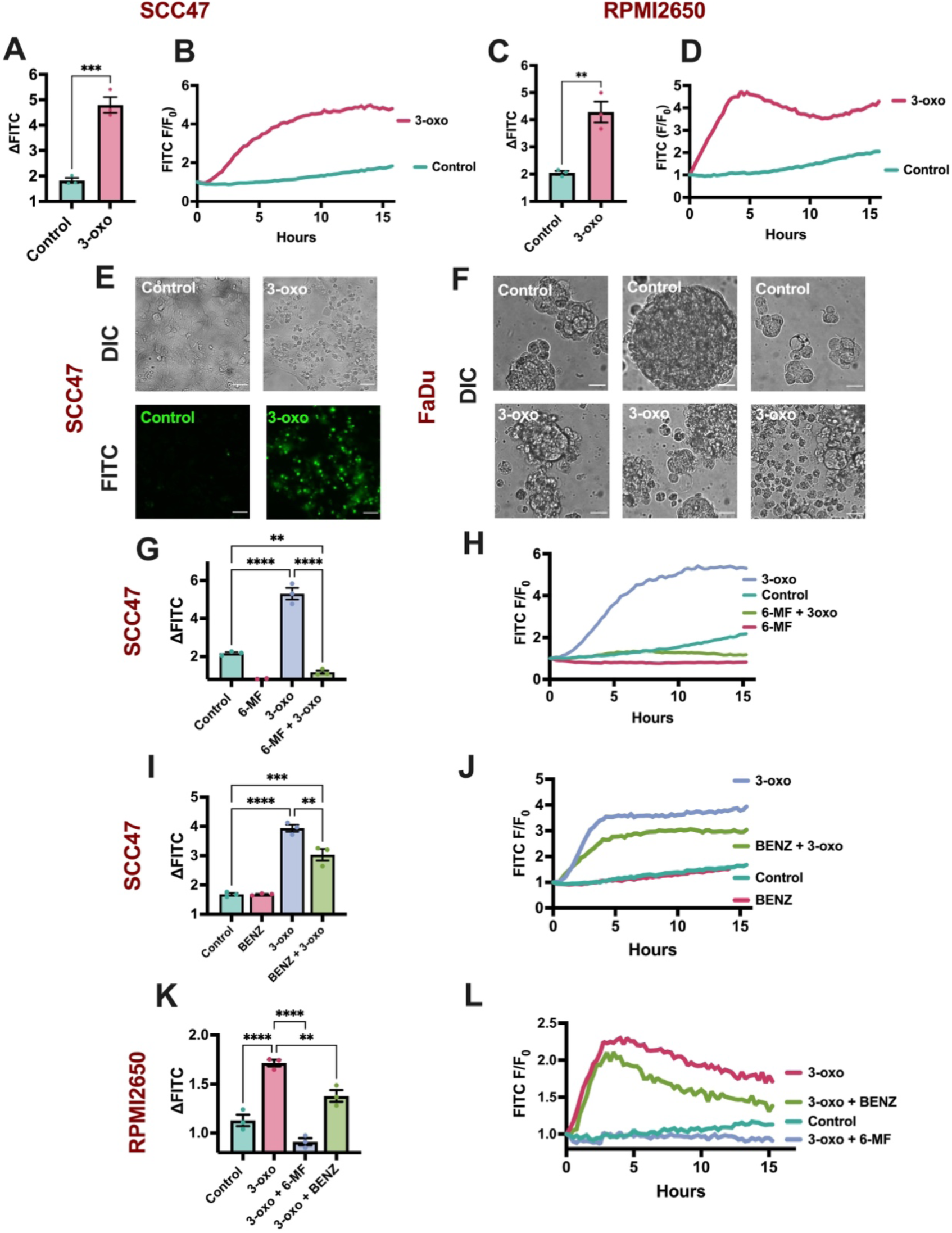
3-oxo-C12HSL induces apoptosis in HNSCC cells via T2R14. Caspase-3 and −7 cleavage and spheroid formation was measured in HNSCC cells. **A & B)** SCC47 Δ CellEvent fluorescence at (*A*) and over (*B*) 15 hours with 100 µM 3-oxo-C12HSL (3-oxo). **C & D)** RPMI2650 Δ CellEvent fluorescence at (*C*) and over (*D*) 15 hours with 100 µM 3-oxo-C12HSL (3-oxo). **E)** Representative images of CellEvent fluorescence and differential interference contrast (DIC) of SCC47 cells after 15 hours with 100 µM 3-oxo-C12HSL (3-oxo). **F)** Representative DIC images of FaDu spheroid formation after 24 hours with 100 µM 3-oxo-C12HSL (3-oxo). **G & H)** SCC47 Δ CellEvent fluorescence at (*G*) and over (*H*) 15 hours with 100 µM 3-oxo-C12HSL (3-oxo) +/- 100 µM 6-MF. **I & J)** SCC47 Δ CellEvent fluorescence at (*I*) and over (*J*) 15 hours with 100 µM 3-oxo-C12HSL (3-oxo) +/- 50 µM benzethonium chloride (BENZ). **K & L)** RPMI2650 Δ CellEvent fluorescence at (*K*) and over (*L*) 15 hours with 100 µM 3-oxo-C12HSL (3-oxo) +/- 50 µM benzethonium chloride (BENZ) or 100 µM 6-MF. All traces are representative. All bar graphs mean ± SEM with >3 experiments using separate cultures. Significance for two conditions was determined with unpaired t-test. Significance for more than two conditions was determined with 1-way ANOVA with multiple comparisons (Control vs 3-oxo; Control vs 3-oxo + 6-MF or BENZ; 3-oxo vs 3-oxo + 6-MF or BENZ) with Šidák correction. All scale bars = 50 µm. (P < 0.05 (*), P < 0.01 (**), P < 0.001 (***), and no statistical significance (ns or unmarked).

### Other Acyl-Homoserine Lactones Do Not Activate T2R14

Many other gram-negative and some gram-positive bacteria produce AHLs [69]. To test if AHLs from other bacterial species activated T2R14 in HNSCCs, commercially available AHLs with carbon chains ranging from 4 to 14 and with and without a ketone or hydroxyl group on the third carbon within the chain (3-oxo or 3-OH) were tested. Strikingly, only 3-oxo-C12HSL induced a Ca^2+^ response in SCC47 and FaDu cells (Figure 8A & B). Using Flamindo 2 to measure cAMP dynamics, 3-oxo-C12HSL increased cAMP while C4-HSL, another *P. aeruginosa* AHL, and 3-oxo-C14HSL did not (Figure 8C & D). In crystal violet assays, both 3-oxo-C12HSL and 3-oxo-C14HSL significantly decreased cell viability in SCC47 and RPMI2650 cells, but only 3-oxo-C14HSL had effects on FaDu cells (Figure 8E-G). Nonetheless, both 3-oxo-C12HSL and 3-oxo-C14HSL inhibited FaDu spheroid formation over 24 hours while C4HSL had no effect.

**Figure 8.**
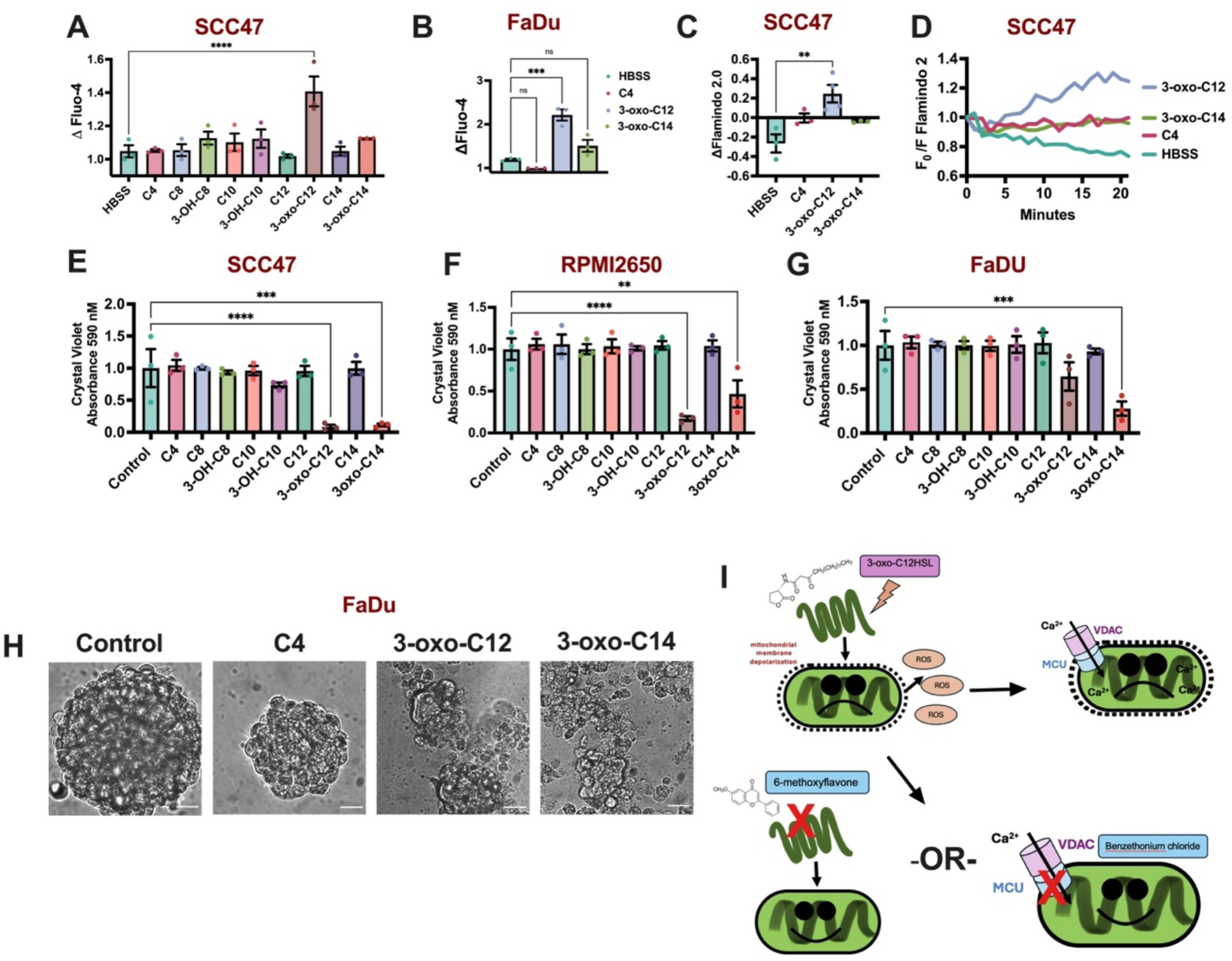
Other acyl-Homoserine Lactones do not activate T2R14. HNSCC cells were loaded with Fluo-4 to measure Ca^2+^ responses. **A)** SCC47 peak Ca^2+^ with 100 µM of each AHL (C4: N-butanoyl-Lhomoserine lactone; C8: N-octanoyl-L-homoserine lactone; 3-OH-C8: N-3-hydroxyoctanoyl-L-homoserine lactone; C10: N-decanoyl-L-homoserine lactone; 3-OH-C10: N-3-hydroxydecanoyl-Lhomoserine lactone; C12: dodecanoyl-L-homoserine lactone; 3-oxo-C12: N-3-oxo-dodecanoyl-homoserine lactone; C14: tetranoyl-L-homoserine lactone; 3-oxo-C14: N-3-oxo-tetranoyl-homoserine lactone). **B)** FaDu peak Ca^2+^ response with 100 µM C4, 3-oxo-C12, or 3-oxo-C14. **C & D)** SCC47 change in cAMP (Δ FITC) (*C*) and response over time (*D*) with 100 µM C4, 3-oxo-C12, or 3-oxo-C14 using Flamindo 2 reporter. **E-G)** Crystal violet assay was used to measure cell viability. SCC47 (*E*), RPMI2650 (*F*), and FaDu (*G*) dissolved crystal violet absorbance values after 24 hours with 100 µM of each AHL. **H)** FaDu spheroid formation over 24 hours with 100 µM C4, 3-oxo-C12, or 3-oxo-C14. **I)** Working model of the mechanism of action of 3-oxo-C12HSL in HNSCC cells. 3-oxo activates T2R14 which causes mitochondrial Ca^2+^ overload via MCU. All bar graphs mean ± SEM with >3 experiments using separate cultures. Significance for more than two conditions was determined with 1-way ANOVA with Dunnett posttest comparing each AHL to HBSS or control. Scale bars = 20 µm. (P < 0.05 (*), P < 0.01 (**), P < 0.001 (***), and no statistical significance (ns or unmarked).

## DISCUSISON

The tumor microenvironment (TME) plays a crucial role in shaping cancer pathogenesis, influencing immune responses, determining treatment outcomes, and affecting biomarker expression [70]. In cancers like head and neck squamous cell carcinoma (HNSCC), which are genetically heterogeneous and challenging to treat, understanding the specific impacts of the TME is essential [71]. The normal oral microbiome can be greatly altered in HNSCCs and thus promote pro-or anti-tumor environments [72]. *P. aeruginosa* exist as a commensal or opportunistic pathogen in the oral microbiome and is more prevalent in HNSCC patients than normal individuals [21, 22, 24]. *P. aeruginosa* secretes 3-oxo-C12HSL, a quorum-sensing molecule acyl homoserine lactone (AHL). 3-oxo-C12HSL is a known apoptosis inducer in several cell types, including cancer, and is a bitter agonist [19, 25, 26]. We found that T2Rs trigger apoptosis in HNSCC cells, with specific activation of T2R14 leading to a strong apoptotic response [9, 10]. While 3-oxo-C12HSL has been shown to activate T2Rs, its interaction with endogenous T2R14 has yet to be investigated. Furthermore, the apoptotic mechanism of 3-oxo-C12HSL has not yet been associated with any T2R, particularly T2R14, and prior research has only focused on the effects of analogues of bacterial AHLs on HNSCC growth [73].

Data here suggest that T2Rs may play chemosensory roles in the HNSCC TME in response to 3-oxo-C12HSL secreted by *P. aeruginosa.* We initially observed a Ca^2+^ response with 3-oxo-C12HSL in several HNSCC cell lines. This Ca^2+^ response was largely linked to endogenous T2R14 activation. However, we did observe differences in Ca^2+^ dynamics in the absence of extracellular Ca^2+^. This may indicate activation of Ca^2+^ channels or receptors other in addition to activation of T2R14. Further supporting this, we observed an SOCE-independent Ca^2+^ response post-ER Ca^2+^ depletion with 3-oxo-C12HSL. Lidocaine, another well documented T2R14 agonist, exhibited dependence on SOCE in our experiments, showing a mechanistic difference in Ca^2+^ signaling between the two T2R14 agonists. Others have suggested that 3-oxo-C12HSL activates TRPV1 [57]. All cell types in this study, except for HEK293T WT, expressed and responded to capsaicin, the exogenous ligand of TRPV1 [59]. Our data support potential activation of TRPV1 by 3-oxo-C12HSL in HNSCC cells. This could be the SOCE-independent mechanism behind the sustained Ca²⁺ responses. Still, 3-oxo-C12HSL activated a largely intact Ca^2+^ response without extracellular Ca^2+^. This is consistent with the observed Ca^2+^ efflux from the ER, dependence on PLC signaling, and the inhibition of the Ca^2+^ with several T2R14 antagonists. Activation of TRPV1 alone with capsaicin was not sufficient to induce apoptosis. While capsaicin slightly decreased cell viability, it was not sufficient for apoptosis induction. This indicates that SOCE and TRPV1 do not underly the apoptotic mechanisms of 3-oxo-C12HSL. While our data suggest that 3-oxo-C12HSL may activate TRPV1 or additional Ca^2+^ channels/receptors, apoptosis with 3-oxo-C12HSL was nonetheless dependent on T2R14 activation. Cell viability was not rescued in ORAI 3KO cells treated with 3-oxo-C12HSL when compared to HEK293T WT cells. Antagonization of T2R14 with 6-MF has a robust rescue effect on apoptosis in the presence of 3-oxo-C12. These results show that T2R14 activation drives apoptosis in HNSCC cells.

T2R14-mediated apoptosis is driven by mitochondrial Ca^2+^ overload, reactive oxygen species (ROS) production, and downstream proteasome inhibition in HNSCC cells [10]. However, the precise mechanism of Ca^2+^ influx into the mitochondria remains unclear. Like other T2R14 bitter agonists, 3-oxo-C12HSL induces both mitochondrial Ca^2+^ overload and ROS generation. We further investigated the specific mitochondrial Ca^2+^ channels that could drive this overload. MCU transports Ca^2+^ into the mitochondria and its function is implicated in many cancers [74]. Mitochondrial Ca^2+^ influx with 3-oxo-C12HSL was diminished when MCU was inhibited. Furthermore, inhibition of MCU rescued cells from mitochondrial depolarization, ROS production, and apoptosis induced by 3-oxo-C12HSL. This reveals, for the first time, a T2R-linked transport mechanism of Ca^2+^ into the mitochondria (Figure 8I). Notably, while 3-oxo-C12HSL has been purported to permeabilize mitochondria, our data suggest a more specific role of MCU in 3-oxo-C12HSL-induced mitochondrial alterations [65].

We observed that 3-oxo-C12HSL elevates cAMP levels in HNSCC cells. Bitter agonists typically decrease cAMP as T2Rs inhibit adenylyl cyclase due to specific coupling to Gα_i_ or Gα_q_ [10]. Previous research demonstrated that 3-oxo-C12HSL enhanced cAMP levels through STIM1 activation in response to ER Ca^2+^ depletion in a lung cancer model. While our results corroborate previous findings, we found that cAMP elevation with 3-oxo-C12HSL is independent from Ca^2+^ signaling and ER-store depletion in HNSCC cells. More work is warranted to elucidate the mechanism behind the 3-oxo-C12HSL-cAMP signaling axis.

Many gram-negative and some gram-positive bacteria employ quorum-sensing AHL molecules to promote biofilm formation and enhance their virulence factors [75]. We investigated if other AHL molecules with long carbon chains activate T2R14 and affect cellular health in HNSCC cells. Only 3-oxo-C12HSL activated significant Ca^2+^ responses and hindered cell viability in HNSCC cells. Interestingly, 3-oxo-C14HSL did decrease cell viability but did not affect Ca^2+^ mobilization. 3-oxo-C14HSL is secreted by *Rhizobium leguminosarum* or *Sinorhizobium meliloti,* which both fix nitrogen and form symbiotic relationships with legumes [76, 77]. Neither of these bacteria are known to infect humans. While 3-oxo-C14HSL could be further investigated as cancer-killing agent, our results suggest that its effects are T2R-independent. C4HSL, which is also produced by *P. aeruginosa*, did not induce a Ca^2+^ response or effect cell viability. This indicates that the anti-cancer effects through AHL secretion in *P. aeruginosa* is specific to 3-oxo-C12HSL.

A recent microbiome analysis of HPV-positive tonsil squamous cell carcinoma revealed that *Burkholderia pseudomallei*, a gram-negative bacteria found in soil, was present in tumor samples [78]. *B. pseudomallei* produce quorum-sensing molecules N-octanoyl-L-Homoserine lactone (C8HSL), N-3-hydroxyoctanoyl-L-Homoserine lactone (3-OH-C8HSL), N-3-hydroxydecanoyl-L-Homoserine lactone (3-OH-C10HSL) [79]. None of these AHLS mobilized Ca^2+^ or affected cell viability. C10HSL, C12HSL, and C14HSL were also tested. These compounds did not have any effects, indicating that from the AHLs tested, 3-oxo-C12HSL alone activates T2R14 in HNSCC cells. This highlights the structural specificity of 3-oxo-C12HSL and the importance of understanding *P. aeruginos*a in the TME of HNSCCs. More work is warranted to explore additional long carbon chain AHLs.

Taken together, we reveal a novel mechanism of 3-oxo-C12HSL-T2R14 activation that may underlie anti-tumor activity of *P. aeruginosa* in the HNSCC TME, and possibly other cancers. *P. aeruginosa* are being explored in the context of cancer treatments by leveraging their apoptotic and immunomodulating properties against cancer [31]. These translational methods could be used to exploit the properties of 3-oxo-C12HSL for HNSCCs. Independent from *P. aeruginosa,* 3-oxo-C12HSL alone could serve as a new bitter agonist therapeutic. Due to the low affinity properties of T2Rs, bitter agonists are often used in the millimolar range for activation [80]. However, 3-oxo-C12HSL is effective in the micromolar range and is commercially available. Beyond cancer, both T2R14 and MCU could serve as therapeutic targets in protecting against apoptosis via *P. aeruginosa* infection in epithelial tissues. Considering the TME in HNSCC is crucial for understanding tumor biology, progression, and treatment responses. Better elucidation of the interactions between T2Rs and endogenous bitter agonists from microbes could lead to more effective therapeutic strategies tailored to the unique features of the HNSCC microenvironment.

## Acknowledgements

This study was supported by a Blavatnik Family Fellowship in Biomedical Research to Z.A.M., a pilot award from the University of Pennsylvania Department of Otorhinolaryngology to R.M.C., and National Institutes of Health grants HL168060 and AI167971 to R.J.L. We thank N. Cohen (University of Pennsylvania) for helpful discussions and M. Victoria (University of Pennsylvania) for technical assistance.

## Author Contributions

Conceptualization, Z.A.M., R.M.C., and R.J.L.; Investigation, Z.A.M., A.M., J.C.T., S.M.S., B.L.H.; visualization, Z.A.M., A.M.; methodology, Z.A.M., J.C.T.; formal analysis, Z.A.M., A.M.; writing – original draft, Z.A.M.; writing – review & editing; Z.A.M., R.M.C., and R.J.L.; funding acquisition, Z.A.M., R.M.C. and R.J.L.; project administration, Z.A.M., R.J.L.; supervision, R.J.L.

## METHODS

### Cell culture

SCC47 (UM-SCC-47; Millipore SCC071), FaDu (ATCC HTB-43), RPMI2650 (ATCC CCL-30), and HEK293T cell lines were from ATCC (Manassas, VA, USA) or MilliporeSigma (St. Louis, MO, USA). ORAI Stable Triple Knockout HEK293 (ORAI 3KO) were from Applied Biological Materials (Ferndale, WA, USA) [81]. All cells were verified by STR sequencing and tested for mycoplasma at least monthly. Cells were grown in high glucose Dulbecco’s modified Eagle’s medium (Corning; Glendale, AZ, USA) with 10% FBS (Genesee Scientific; El Cajon, CA, USA), penicillin/streptomycin mix (Gibco; Gaithersburg, MD, USA), and nonessential amino acids (Gibco). 0.25% or 0.05% EDTA Trypsin (Gibco, Gaithersburg, MD, USA) was used at 37 °C for 10 min for single-cell suspension solution for experimental plating. Cells were utilized in experiments when cell confluency in culture vessel reached 70–80%. Cell lines were not utilized past passage 25.

### Pharmacological Agents and Stock Solutions

*N*-(3-Oxododecanoyl)-L-homoserine lactone (3-oxo-C12HSL) (MilliporeSigma; St. Louis, MO, USA was dissolved in dimethyl sulfoxide (DMSO) at 100 mM. All subsequent experiments used 3-oxo-C12HSL at 100 μM or less, resulting in ≤ 0.1% DMSO in all experimental samples. All other AHLs (Cayman Chemical; Ann Arbor, MI, US) were used dissolved and used in the same way. 3-oxo-C12HSL sourced from Cayman Chemical was also used in experiments to validate effects from different suppliers. Unless specified, all other pharmacological agents were dissolved in water or DMSO at 1000x concentration and used 1:1000, resulting in ≤ 0.1% DMSO in each condition..

### Quantitative reverse transcription PCR (qPCR)

Cells were harvested and lysed in TRIzol when culture reached 70% confluency for measures of endogenous expression. RNA was isolated and purified using product protocol (Direct-zol RNA kit; Zymo Research). Final RNA was eluted in 25 - 30 μL of nuclease free water. RNA concentration of samples was measured using Tecan (Spark 10M; Mannedorf, Switzerland) NanoQuant plate. Between 200 – 500 ng of RNA were used to transcribe cDNA using MultiScribe Reverse Transcriptase (High-Capacity cDNA Kit; Themo Fisher Scientific). A PCR Thermocycler was used to synthesize cDNA in one cycle of 25 °C for 10 minutes, 37 °C for 120 minutes, and 85 °C for 5 minutes. Samples were diluted with nuclease free water to 5 ng cDNA per μL (assuming ng of RNA in rxn = ng of cDNA synthesized). Gene expression in cDNA samples were quantified using TaqMan qPCR probes for *TAS2R14, TRPV1, TRPA1, VDAC1, VDAC2, VDAC3, MCU, SLC8B1,* and *UBC* (Thermo Fisher Scientific). 2.5 μL of Fast Advanced, 0.25 μL of TaqMan qPCR probes, and 0.25 μL of Nuclease free water was used per well (384-well qPCR plate) in combination of 2 μL of cDNA (at 5 ng/μL). This was done in triplicate for each sample. Gene expression was measure using QuantStudio 5 Real-Time PCR System for Human Identification during 40 cycles of 50 °C for 2 minutes, 95 °C for 2 minutes, 95 °C for 1 second, 60 °C for 20 seconds. Gene expression was calculate using *UBC* as an endogenous control due to stable expression in cancer cells (2).

### Live Cell Imaging

For calcium (Ca^2+^) imaging, cells were loaded with 5 µM of Fluo-4-AM (Thermo Fisher Scientific) or for 50 min at room temperature in the dark. Hank’s Balanced Salt Solution (HBSS) buffered with 20 mM HEPES (pH 7.4) was used as an imaging buffer, containing 1.8 mM Ca^2+^. Cells were imaged using an Olympus IX-83 microscope (20x 0.75 NA PlanApo objective), FITC filters (Semrock FF01-474/27 Ex; Semrock FF01-525/45 Em) in excitation and emission filter wheels (Sutter Lambda LS), Orca Flash 4.0 sCMOS camera (Hamamatsu, Tokyo, Japan), Meta-Fluor (Molecular Devices, Sunnyvale, CA USA), and XCite 120 LED Boost (Excelitas Technologies).

For experiments in Ca^2+^-free (0-Ca^2+^) HBSS, cells were loaded with Fluo-4-AM as described above in normal Ca^2+^-containing HBSS. Agonists were dissolved in HBSS with no added Ca^2+^ plus 2 mM EGTA (0-Ca^2+^ HBSS) and was used in the place of regular HBSS (no EGTA). At the start of the experiment, cells were washed 1 x in 0-Ca^2+^ HBSS and left in 30 – 100 µL 0-Ca^2+^ HBSS. For experiments in which Ca^2+^ was added back after initial stimulation, HBSS containing 8 mM Ca^2+^ was added post-initial response. With the addition of 10 mM Ca^2+^ into 2 mM EGTA HBSS, predicted free extracellular Ca^2+^ was 2 mM (MaxChelator, Chris Patton, Stanford University, https://somapp.ucdmc.ucdavis.edu/pharmacology/bers/maxchelator/).

For mitochondrial and endoplasmic reticulum Ca^2+^ imaging, pcDNA-4mt3cp, pcDNA-D1ER, or CMV-ER-LAR-GECO1.2 transfected into cells using lipofectamine 3000 reagents 24-48 hours prior to imaging.[38, 39] Live cell images were taken on Olympus IX-83 microscope (20x 0.75 NA PlanApo objective), CFP/YFP (Chroma ET430/24x Ex; Chroma ET525/30m Em) (pcDNA-4mt3cp and pcDNA-D1ER) or TRITC (Semrock FF01-554/23 Ex; Semrock FF01-609/54 Em) (CMV-ER-LAR-GECO1.2) filters in excitation and emission filter wheels (Sutter Lambda LS), Orca Flash 4.0 sCMOS camera (Hamamatsu, Tokyo, Japan), Meta-Fluor (Molecular Devices, Sunnyvale, CA USA), and XCite 120 LED Boost (Excelitas Technologies). For cyclic adenosine monophosphate (cAMP) imaging, pcDNA3.1(-)Flamindo2 was transfected into cells as described above 48 hours prior to imaging.[47] Cells were imaged as described above using FITC filters. pcDNA3 pAMPKar and Sapphire AKAR was used to measure AMPK or PKA activity, respectively, using microscope and CFP/YFP filters as described above [49, 82]. To measure specific nuclear or cytoplasmic Ca^2+^, pCMV G-GECO1.2 or pCMV NLS-R-GECO were used and imaged on microscope and TRITC filters described above [37].

For Ca^2+^ imaging using supernatant from *Pseudomonas aeruginosa* strain PAO1 or JP2, single colonies were selected and grown in LB broth for 72 hours until PAO1 cultures turned green, indicating 3-oxo-C12HSL production [83, 84]. Cultures were centrifuged and supernatant was collected. It was determined that HBSS containing ≤ 12.5% LB did not have a significant Ca^2+^ response, thus supernatants were diluted down to 25% in HBSS for stimulation (12.5% when added to chamber slide culture). Fluo-4 was used as the Ca^2+^ indicator dye.

### Cell Viability

Crystal Violet assay was to measure cell viability. Cells were treated for 6, 24, or 48 hours +/- 0, 1, 10, or 100 µM 3-oxo-C12HSL in DMEM high-glucose. Treatment was removed following incubation time and cells were washed 1x with PBS. Crystal violet 0.1% in deionized water was used to stain remaining adherent, viable cells (200 μL per well in 48 well plate). Crystal violet stayed on cells for 5 minutes for uptake of stain. Crystal violet was aspirated, and remaining stains were washed 3x with deionized water and left to dry for 24 hours. Crystal violet stains were then dissolved with 30% acetic acid in deionized water. Absorbance at 590 nm measured on a Tecan (Spark 10M; Mannedorf, Switzerland). Absorbance values for experimental wells were normalized to control (no treatment).

XTT assay was done according to measure NADH production. 30 μL of XTT dye (stock 0.1 mg/µL in DMSO) and 7.5 μL phenazine methosulfate (stock 3 mg/mL in PBS) (Millipore Sigma) were added to 3 mL of phenol-free DMEM. Cells (in 48 well plate) were washed 3x with PBS and treated at 1.25x normal concentration of 3-oxo-C12HSL ranging from 0 – 100 µM 3-oxo-C12HSL in phenol-free DMEM (200 µL per well). 50 μL of XTT/phenazine methosulfate solution was added to each well (adjusting final treatment concentration to 1x). Absorbance values were measured at 475 and 660 nm every 2 hours for 6 hours on Tecan Spark 10M. Final values were calculated by subtracting 660 nm wavelength from 475 nm wavelength and normalizing to control well(s).

### Spheroid Formation

FaDu cells were trypsonized into single-cell suspension and plated on low-attachment 96-well plate in DMEM/F-12 1:1 media supplemented with 10 ng/μL epidermal growth factor (EGF) and fibroblast growth factor (FGF) and 1% B-17 Supplemental [85]. HSL molecules (C4HSL, 3-oxo-C12HSL, or 3-oxo-C14HSL) were added to each culture while plating. After 24 hours, live cell DIC images were taken on Olympus IX-83 microscope system to observe spheroid formation.

### Mitochondrial Health

A mitochondrial membrane potential indicator dye, TMRE, was loaded into cells on a glass-bottom at 500 nM in phenol-free DMEM [68]. Cells were loaded for 15 minutes in the dark at RT. Dye was aspirated and cells were washed 3x with PBS. +/- 3-oxo treatment in phenol-free DMEM was added to each appropriate well. Fluorescence was measured on Tecan Spark 10 M on FITC and TRITC filters every 15 minutes over 15 hours at 37 °C, 5% CO2. TRITC fluorescence intensity was normalized to time 0 and plotted over time. To measure mitochondrial superoxide (ROS) species, MitoSox dye was added to cells at 500 nM in phenol-free DMEM in parallel with +/- 3-oxo treatment. Fluorescence was measured as described above. Representative images taken with TRTIC filter on Olympus IX-83 microscope using 20x objective at end of 15-hour assay period. Fluorescence intensity adjusted using the HiLo look-up table (LUT) to best show observed results. Intensity values were set equally across all images compared or normalized to control for quantification.

### CellEvent

CellEvent Green Caspase 3/7 dye (ThermoFisher Scientific) was added to cells in a glass bottom plate with +/- 3-oxo in phenol-free DMEM. For every 500 μL of phenol-free DMEM treatment, 1 drop of CellEvent reagent was used (dropper bottle). Quantitative fluorescence values measured on a Tecan Spark 10M over 15 hours at 37 °C, 5% CO2. Representative images taken with FITC filter on Olympus IX-83 microscope using 20x objective after 15 hours. Fluorescence intensity adjusted using the HiLo look-up table (LUT) to best show observed results. Intensity values were set equally across all images compared or normalized to control for quantification.

### Data analyses and statistics

Raw data points in all bar graphs represents ≥ 3 independent experiments with center representing the mean and error bars representing SEM. Data were analyzed using *t-*test (two comparisons only), one-way ANOVA (>2 comparisons), two-way ANOVA (>1 group), or area under curve in GraphPad Prism (San Diego, CA, USA). Both paired and unpaired *t*-test were used when appropriate. Dunnett or Bonferroni posttest or Šidák correction was used for one-way ANOVA, or two-way ANOVA were used when necessary. In all figures, <0.05 (*), <0.01(**), and <0.001(***) while no statistical difference was unlabeled.

