## Supplementary Figures for "*Pseudomonas aeruginosa* metabolite 3-oxo-C12HSL induces apoptosis through T2R14 and the mitochondrial calcium uniporter"

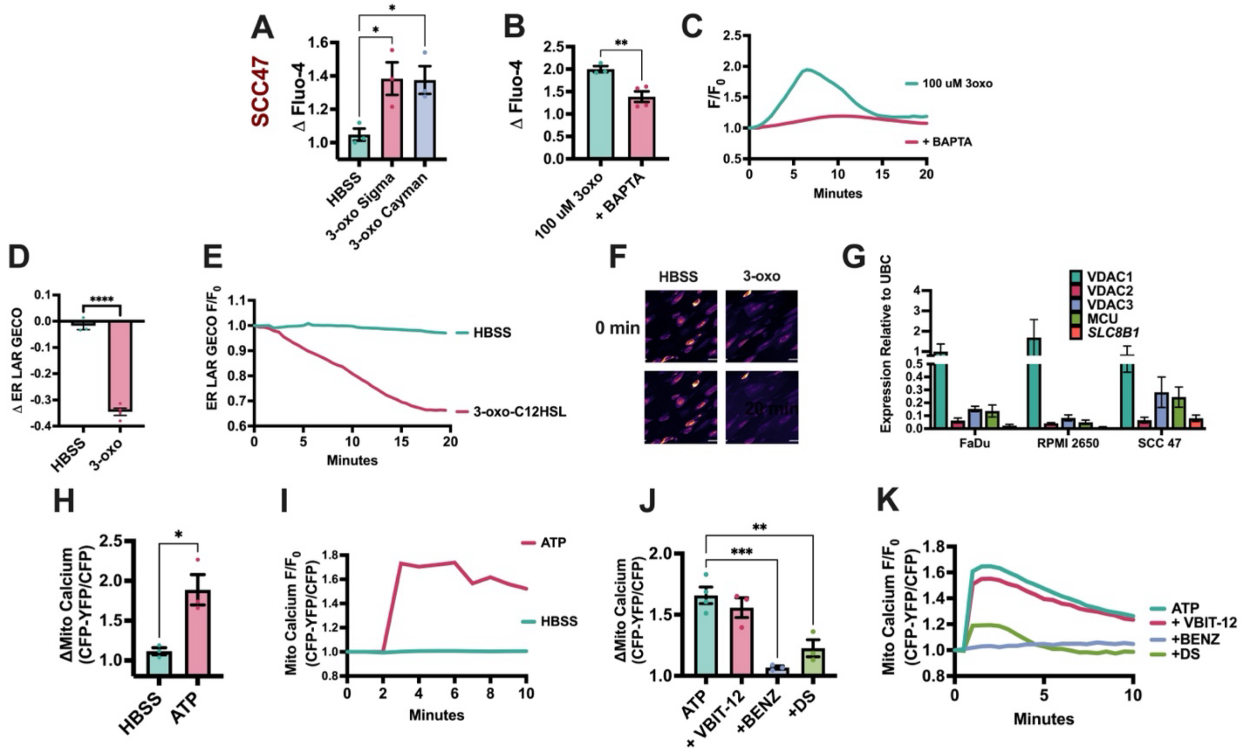


**Supplemental Figure 1. 3-oxo-C12HSL activates an intracellular $Ca^{2+}$ response.** HNSCC cell line SCC47 was loaded with Fluo-4 or transfected with a genetically encoded $Ca^{2+}$ reporter. **A)** SCC47 peak $Ca^{2+}$ responses with 100 μM 3-oxo-C12HSL from Sigma (MilliporeSigma; St. Louis, MO USA) or Cayman (Cayman Chemical; Ann Arbor, MI, US). **B & C)** Peak $Ca^{2+}$ response (*B*) and response over time (*C*) with 100 μM 3-oxo-C12HSL loaded with +/- 10 μM BAPTA-AM. **D-F)** ER $Ca^{2+}$ efflux change (Δ) (*D*) and over time (*E*) in CFP-YFP/CFP fluorescence with 100 μM 3-oxo-C12HSL (3-oxo) using ER LAR GECO reporter (*F*) (scale bar = 50 μm). **G)** mRNA expression of *VDAC1, VDAC2, VDAC3, MCU, and SLC8B1* in FaDu, SCC47, and RPMI2650 cells. **H & I)** Mitochondrial $Ca^{2+}$ influx change (Δ) (*H*) and over time (*I*) in CFP-YFP/CFP fluorescence with 100 μM 3-oxo-C12HSL (3-oxo) using 4mt3Dcpv mitochondrial $Ca^{2+}$ reporter. **J & K)** Mitochondrial $Ca^{2+}$ influx change (Δ) (*H*) and over time (*I*) in CFP-YFP/CFP fluorescence with 100 μM 3-oxo-C12HSL (3-oxo) +/- using 4mt3Dcpv mitochondrial $Ca^{2+}$ reporter +/- 50 μM VBIT-12 (VBIT-12), 50 μM DS16570511 (DS), or 50 μM benzethonium chloride (BENZ). All traces are representative. All peak $Ca^{2+}$ responses mean ± SEM with >3 experiments using separate cultures. Significance for two conditions was determined with unpaired t-test. Significance 1-way ANOVA with Bonferroni’s posttest comparing HBSS/Control to each condition. P < 0.05 (*), P < 0.01 (**), P < 0.001 (***), and no statistical significance (ns or unmarked).

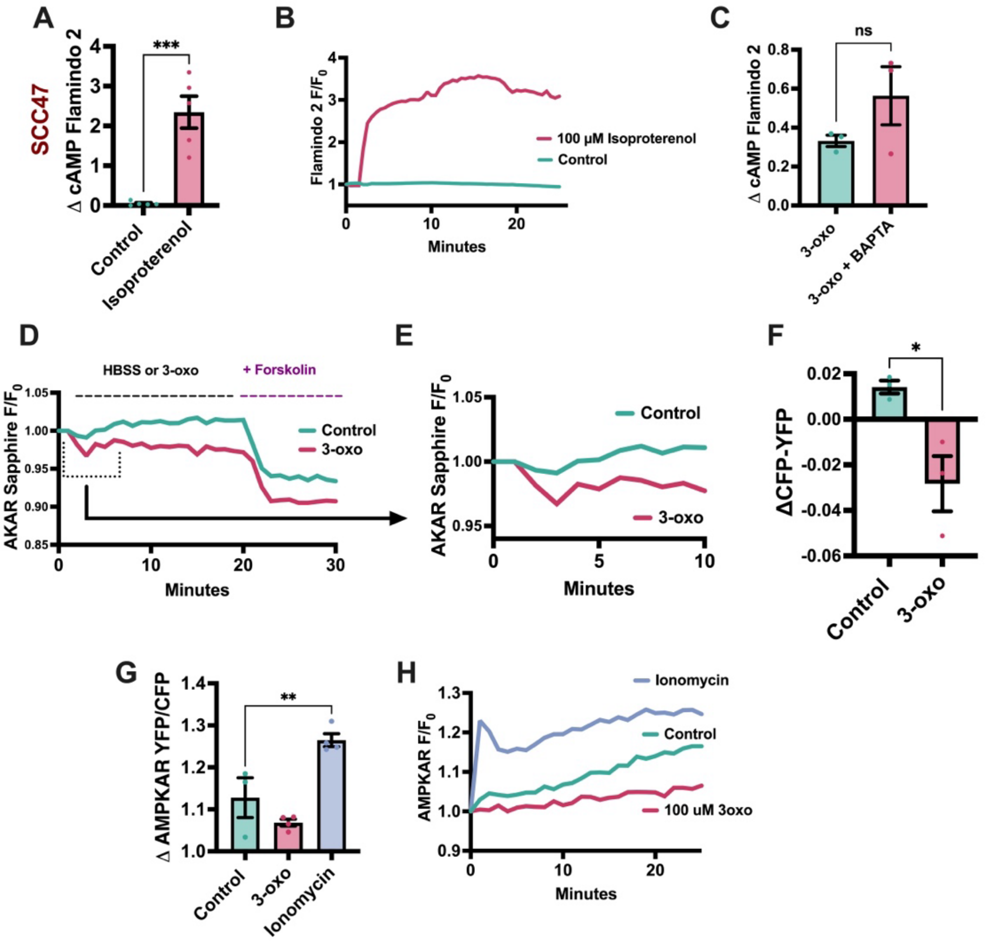


**Supplemental Figure 2. 3-oxo-C12HSL modulates cAMP levels.** cAMP dynamics were measured with SCC47 cells using genetically encoded cAMP, PKA, or AMPK reporters. **A & B)** cAMP change (Δ) *(A)* and response over time (*B*) in FITC with 100 μM isoproterenol using Flamindo 2 cAMP reporter. **C)** cAMP change (Δ) in FITC with 100 μM 3-oxo-C12HSL (3-oxo) loaded with +/- 10 μM BAPTA-AM using Flamindo 2 cAMP reporter. **D-F)** PKA response over time (*D & E*) and change (Δ) (*F*) in CFP-YFP using Sapphire-AKAR PKA reporter. **G & H)** AMPK activation change (Δ) (G) and AMPK response over time (H) in YFP/CFP using AMPKAR AMPK reporter. All traces are representative. All peak $Ca^{2+}$ responses mean ± SEM with >3 experiments using separate cultures. Significance for two conditions was determined with unpaired t-test. Significance 1-way ANOVA with Bonferroni's posttest comparing HBSS/Control to condition. $P < 0.05$ (*), $P < 0.01$ (**), $P < 0.001$ (***), and no statistical significance (ns or unmarked).

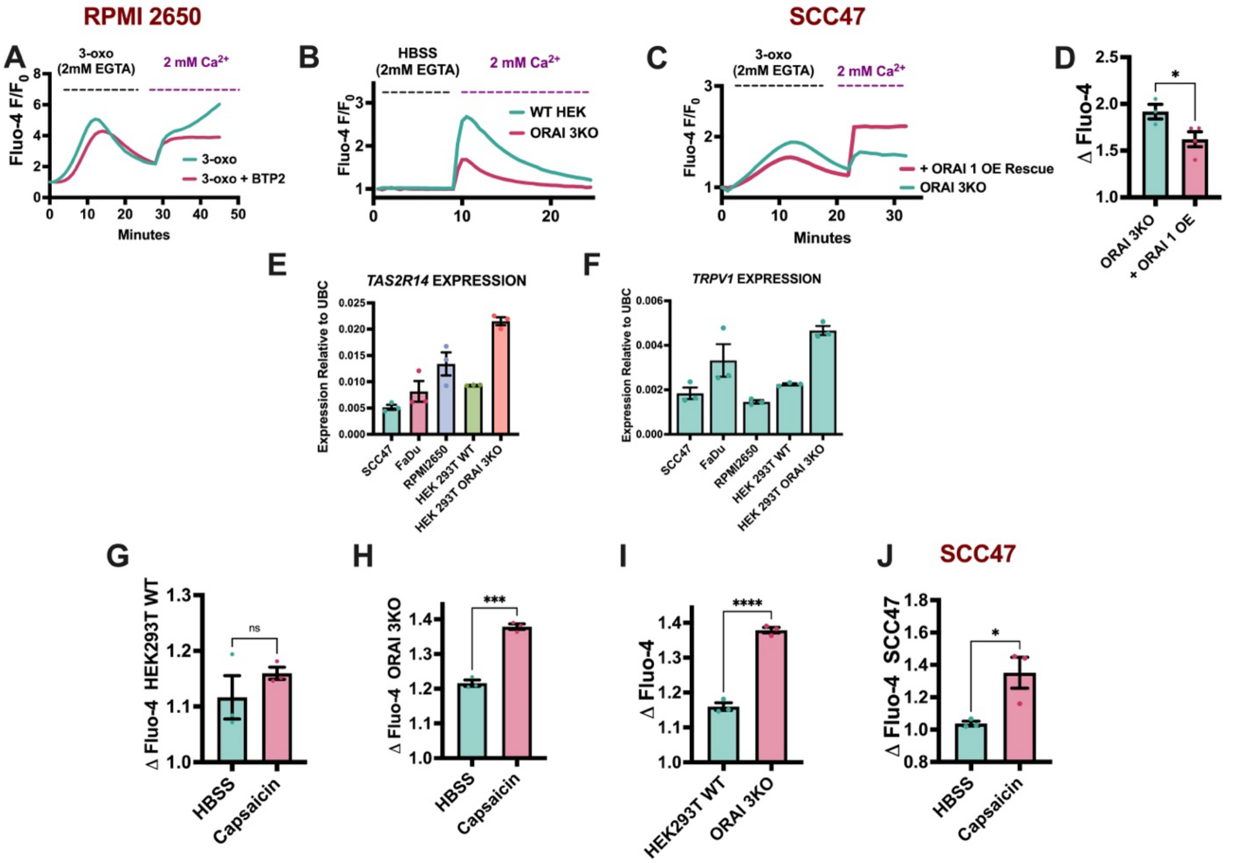


**Supplemental Figure 3. Sustained 3-oxo-C12HSL $Ca^{2+}$ mobilization may be dependent on TRPV1 activation. A)** RPMI2650 $Ca^{2+}$ trace to first stimulation with 100 μM 3-oxo-C12SHL (3-oxo) in 2 mM EGTA and subsequent stimulation with 2 mM $Ca^{2+}$ addback +/- 10 μM BTP2. **B)** HEK293T WT and ORAI 3KO $Ca^{2+}$ trace first stimulation with HBSS in 2 mM EGTA and subsequent stimulation with 2 mM $Ca^{2+}$ addback. **C & D)** ORAI 3KO HEK293 and ORAI 3KO + ORAI1 OE rescue $Ca^{2+}$ trace (*C*) and peak $Ca^{2+}$ response (*D*) to first stimulation with 100 μM 3-oxo-C12SHL (3-oxo) in 2 mM EGTA and subsequent stimulation with 2 mM $Ca^{2+}$ addback. **E & F)** mRNA expression of *TAS2R14* (*E*) and *TRPV1* (*F*) in SCC47, FaDu, RPMI2650, HEK293T WT, and ORAI 3KO HEK293 cells. **G-J)** Peak $Ca^{2+}$ responses in HEK293T WT (G) and ORAI 3KO HEK293 (H) with 100 μM capsaicin. Peak responses between HEK293T WT and ORAI 3KO HEK293 cells were compared (*I*). Peak $Ca^{2+}$ in SCC47 cells with 100 μM capsaicin (*J*). Significance for two conditions was determined with unpaired t-test. P < 0.05 (*), P < 0.01 (**), P < 0.001 (***), and no statistical significance (ns or unmarked).

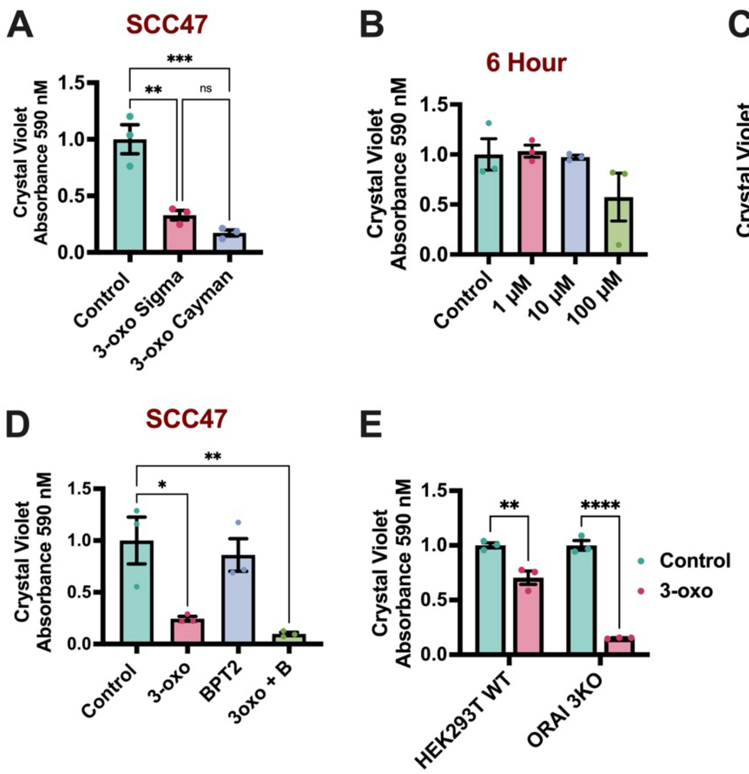


**Supplemental Figure 4. 3-oxo-C12HSL decreases cell viability independent of TRPV1 and SOCE.** Cells were stained with crystal violet, indicating alive/adherent cells post-treatment in normal media conditions. Solubilized crystal violet absorbance was measured at 590 nM and quantified**. A)** SCC47 cells were treated with 100 μM 3-oxo-C12HSL from Sigma (MilliporeSigma; St. Louis, MO USA) or Cayman (Cayman Chemical; Ann Arbor, MI, US) for 24 hours. **B & C)** SCC47 cells were treated with 0 – 100 μM 3-oxo-C12HSL for 6 (*B*) or 48 (*C*) hours. **D)** SCC47 cells were treated with 100 μM 3-oxo-C12HSL +/- 10 μM BTP2 for 24 hours. **E)** HEK293T WT and ORAI 3KO cells were treated with 100 μM 3-oxo-C12HSL for 24 hours. **F)** SCC47 cells were treated with 100 μM 3-oxo-C12HSL or capsaicin for 24 hours. All bar graphs ± SEM with >3 experiments using separate cultures. Significance by 1-way ANOVA with Bonferroni’s posttest comparing HBSS/Control to each dose response or with multiple comparisons with Šidák correction. Significance by 2-way ANOVA for panel *E* with Šidák correction. P < 0.05 (*), P < 0.01 (**), P < 0.001 (***), and no statistical significance (ns or unmarked).

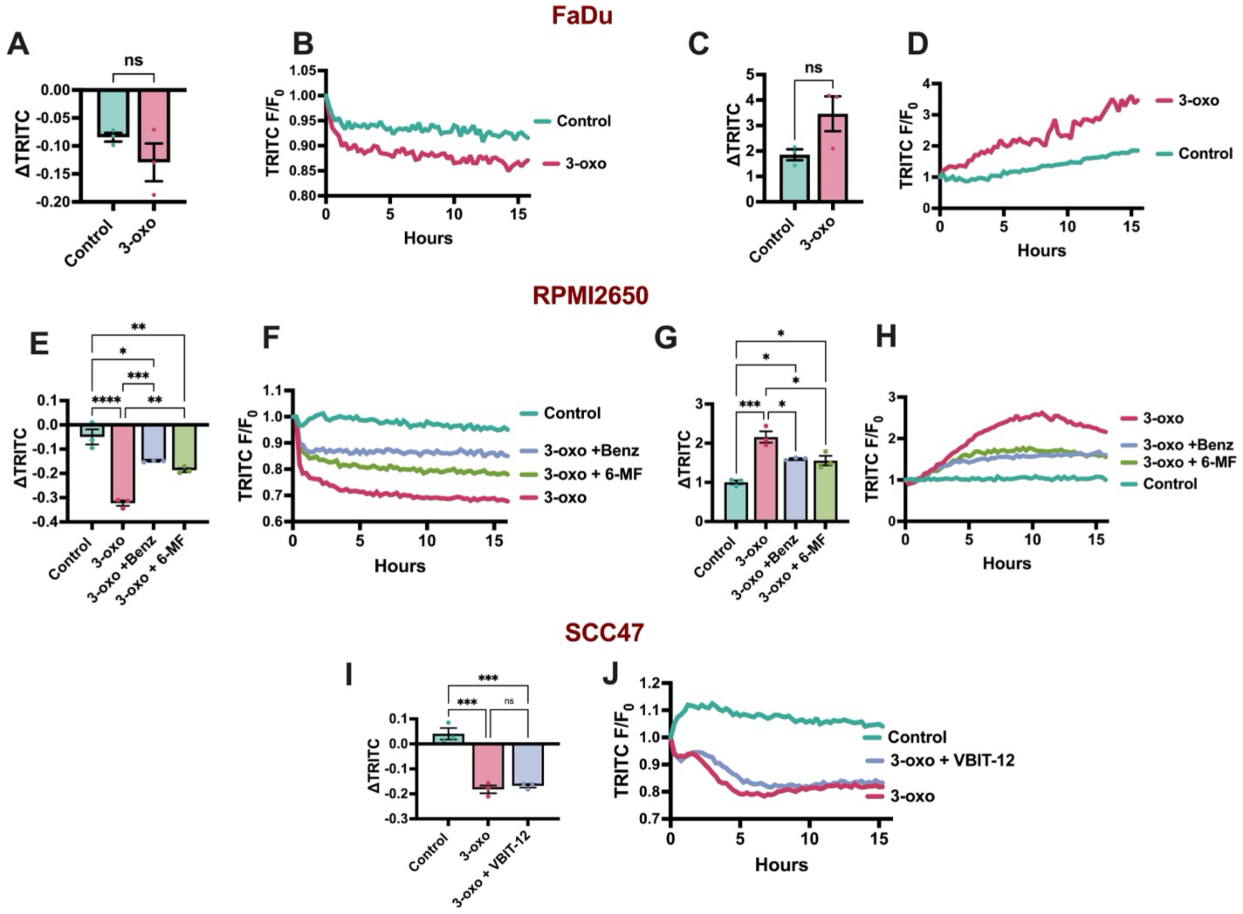


**Supplemental Figure 5. 3-oxo-C12SHL depolarizes mitochondrial membrane potential and produces ROS.** Mitochondrial health and ROS production was measured in FaDu, RPMI2650 and SCC47 cells. **A-D)** FaDu ΔTMRE fluorescence (TRITC) after 15 hours (*A*) and over time (*B*) with 100 μM 3-oxo-C12HSL (3-oxo). FaDu ΔMitoSox fluorescence (TRITC) after 15 hours (*C*) and over time (*D*) with 100 μM 3-oxo-C12HSL (3-oxo). **E-H)** RPMI2650 ΔTMRE fluorescence (TRITC) after 15 hours (*E*) and over time (*F*) with 100 μM 3-oxo-C12HSL (3-oxo) +/- 50 μM benzethonium chloride (BENZ) or 100 μM 6-methoxyflavone (6-MF). RPMI2650 ΔMitoSox fluorescence (TRITC) after 15 hours (*G*) and over time (*H*) with 100 μM 3-oxo-C12HSL (3-oxo) +/- 50 μM benzethonium chloride (BENZ) or 100 μM 6-methoxyflavone (6-MF). **I & J)** SCC47 ΔTMRE fluorescence (TRITC) after 15 hours (*I*) and over time (*J*) with 100 μM 3-oxo-C12HSL (3-oxo) +/- 50 μM VBIT-12. All traces are representative. All bar graphs mean ± SEM with >3 experiments using separate cultures. Significance for two conditions determined with unpaired t-test. Significance for more than two conditions was determined with 1-way ANOVA with multiple comparisons (Control vs 3-oxo; Control vs 3-oxo + 6-MF or BENZ; 3-oxo vs 3-oxo + 6-MF or BENZ or VBIT-12) with Šidák correction. (P < 0.05 (*), P < 0.01 (**), P < 0.001 (***), and no statistical significance (ns or unmarked).

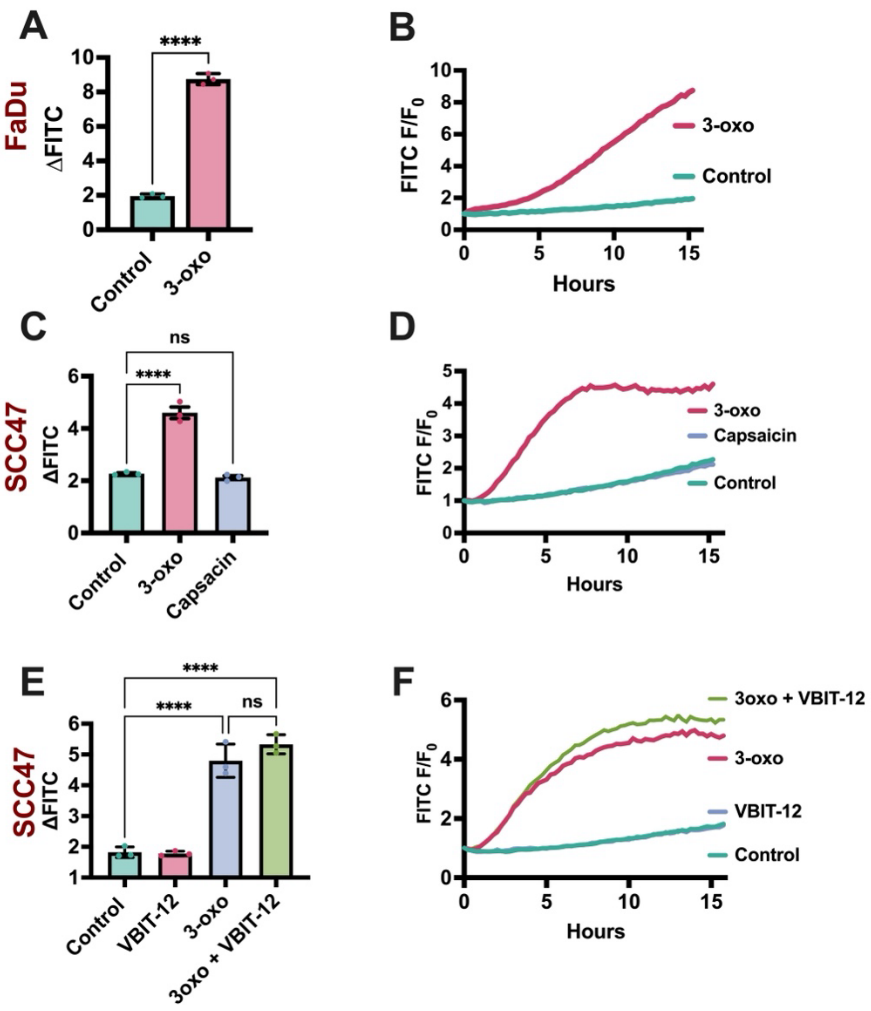


**Supplemental Figure 6. 3-oxo-C12HSL induces apoptosis in HNSCC cells.** Caspase-3 and -7 cleavage was measured in HNSCC cells. **A & B)** FaDu Δ CellEvent fluorescence at (*A*) and over (*B*) 15 hours with 100 µM 3-oxo-C12HSL (3-oxo). **C & D)** SCC47 Δ CellEvent fluorescence at (*A*) and over (*B*) 15 hours with 100 µM 3-oxo-C12HSL (3-oxo) or 100 µM capsaicin. **E & F)** SCC47 Δ CellEvent fluorescence at (*A*) and over (*B*) 15 hours with 100 µM 3-oxo-C12HSL (3-oxo) +/- 50 µM VBIT-12. Significance for two conditions determined with unpaired t-test. Significance for more than two conditions was determined with 1-way ANOVA comparing treatment to control or with multiple comparisons (Control vs 3-oxo; Control vs 3-oxo + VBIT-12; 3-oxo vs 3-oxo + VBIT-12) with Šidák correction. (P < 0.05 (*), P < 0.01 (**), P < 0.001 (***), and no statistical significance (ns or unmarked).
